# Integrative Biophysical Illumination of the 3D GPCRome Dynamics

**DOI:** 10.1101/2025.10.27.684741

**Authors:** Brian Medel-Lacruz, David Aranda-García, Aditya Prasad Patra, Tomasz Maciej Stepniewski, Adrian García-Recio, Mariona Torrens-Fontanals, Franz Hagn, Massimiliano Bonomi, Jiafei Mao, Jana Selent

## Abstract

Achieving high-resolution insights into protein dynamics remains a major challenge, particularly for membrane proteins such as G protein–coupled receptors (GPCRs) - a highly dynamic class of cell surface receptors and key drug targets. Here, we present an integrative strategy that bridges computational and experimental biophysical methods for probing protein dynamics. To illustrate its potential, we developed GPCRmd-NMR, a platform that synergistically combines molecular dynamics (MD) simulations with high-resolution nuclear magnetic resonance (NMR) spectroscopy across the 3D GPCRome. The resulting dataset maps over 20 billion NMR signals to more than 7 million conformational snapshots of GPCRs, enabling structure-resolved analysis at an unprecedented scale. Application across multiple GPCRs provides access to previously underexplored functional GPCR dynamics, receptor states, and sites, including microswitches and hidden ligand entrance gateways. Our integrated framework offers a powerful solution to long-standing challenges faced by both experimental and computational researchers. To support broad accessibility, GPCRmd-NMR is available as an open-access, web-based framework equipped with user-friendly analysis tools (https://www.gpcrmd.org/gpcrmd-nmr-tool) and is readily applicable to other dynamic protein families beyond GPCRs and other types of biophysical data.

## Introduction

G protein-coupled receptors (GPCRs) are an important class of cell surface receptors that sense a large variety of extracellular stimuli such as light, bitter taste, odor, hormones, neurotransmitters, and mechanical stresses. The crucial roles of GPCRs in numerous physiological and pathological processes have made them popular drug targets^1^. Therefore, the molecular mechanisms of GPCR functions and regulations present a central scientific question in both academia and the health industry. During the last two decades, X-ray crystallography and, more recently, cryo-electron microscopy (cryo-EM) have provided valuable high-resolution insights into the distinct functional states of these proteins^2,3^. However, the protein structure data provided by these techniques only present certain particularly stable conformations, leaving many other potentially functional and druggable conformations of these inherently dynamic receptors underexplored. To access these “dark” conformational spaces, spectroscopic and computational methods have been playing leading roles in mapping the structural dynamics governing GPCR signaling^4^.

Spectroscopic methods, including nuclear magnetic resonance (NMR)^5^, electron paramagnetic resonance (EPR)^6^, single-molecule fluorescence resonance energy transfer (smFRET)^5^, and other optical spectroscopic approaches (e.g., bimane-based fluorescence spectroscopy) map the structural changes of receptors at the atomic to nanometer scales via strategically placed reporters. Computational approaches, represented by molecular dynamics (MD) simulations, depict the structural evolution of proteins at the atomistic resolution in a full *in silico* setting. Both methods have provided unique structural information on ligand binding^7^, ligand-induced receptor activation^8,9^, recognition of intracellular coupling partners^10–12^ or lipid/cholesterol interactions^13,14^. Despite the great success of these methods in resolving GPCR dynamics and dynamics-governed signaling, major gaps remain in both method domains. The spectroscopic methods are restricted to a limited number of detectable sites as well as the types of readouts, hampering the comprehensive high-resolution mapping of GPCR dynamics. Additionally, practical challenges — including the selection of suitable reporters and readouts, data interpretation, and extrapolation of results to full receptor dynamics — have restricted the accessibility of these powerful techniques. MD simulations, on the other hand, while providing a fully atomistic view of proteins, are highly dependent on expertise and computational resources to reach meaningful timescales.

Over the past years, we have tackled the challenges associated with MD simulations by the development of a community-based MD approach, namely GPCRmd, which has democratized the MD simulations in the GPCR and related fields via a low-barrier, self-customizable and resource-shared open-access platform^15,16^. This landmark has paved the way for accessing receptor dynamics by integrating computational and spectroscopic approaches with unprecedented spatiotemporal resolution. Here, we showcase the power of this approach on GPCRs with a widely used, high-resolution spectroscopic technique, namely NMR^17–19^ in both solution and solid-state. Our GPCRmd-NMR platform streams *in-silico* MD simulation to spectroscopic features, unveiling previously hidden structural dynamics of GPCRs across functional states. It unlocks the potential of spectroscopic and computational biophysical methods in depicting and deciphering functionally relevant dynamics at the 3D GPCRome level with over 20 billion NMR signals assigned to more than 7 million GPCR conformations.

Our work represents an innovative framework that can be expanded feasibly to other spectroscopic and computational methods, presenting a gateway to data science-driven biophysical and structural biology method integration. Furthermore, it forms the foundation for developing new machine-learning approaches for biophysics and protein science. Ultimately, the established GPCRmd-NMR platform can also serve as a template for large-scale integrative interrogation of function-relevant protein dynamics of other important protein families (e.g. kinome).

## Results

### Chemical shift streaming for the dynamic 3D GPCRome

We have focused on the chemical shift (CS), a site-specific NMR parameter that is highly sensitive to protein structure, molecular interaction and local environment. In the endeavor of forward computational prediction of CSs from protein structures, a number of high-performance predictors have been developed^20^. Two representative and widely used predictors are SHIFTX2^21^ and SPARTA+^22^. These predictors primarily rely on machine learning (ML) and are trained on experimental protein NMR data, capturing the sophisticated relationship between protein structures and chemical shifts. Taking these well-established chemical shift predictors, we have integrated the in-frame chemical shift prediction with MD simulations (**Figure 1A**) across the 3D GPCRome.

**Figure 1.**
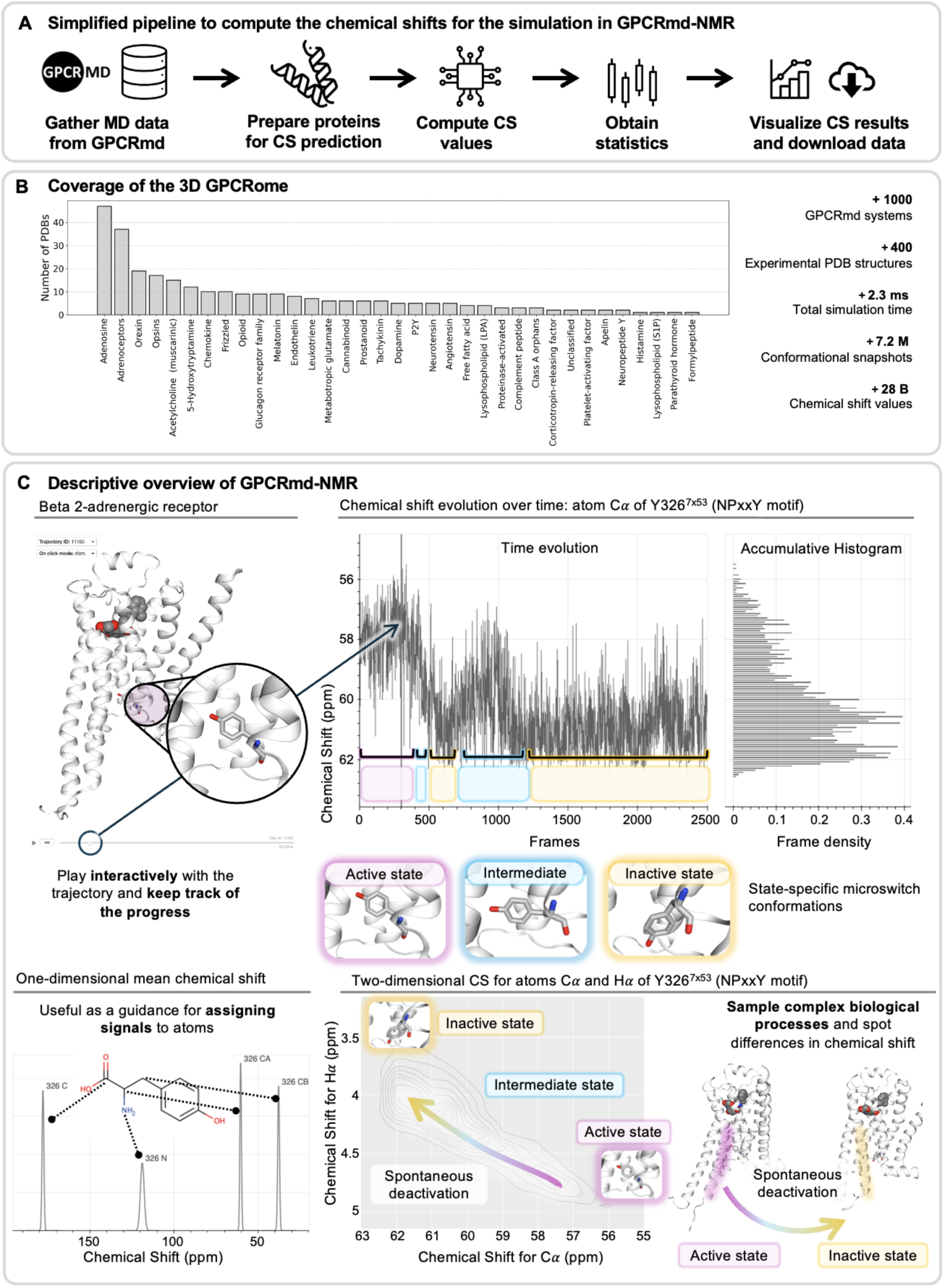
Streaming chemical shifts for GPCRs. (**A**) A simplified pipeline used to compute the chemical shifts for the simulations in GPCRmd-NMR. All data is downloadable as CSV files. One CSV file includes the predicted chemical shift for each frame across all atoms, while a second CSV file (condensed format) provides the mean value, deviation, and an estimation of the prediction-associated error for all frames of the MD. (**B**) Coverage of the 3D GPCRome. **(C**) Descriptive overview of GPCRmd-NMR using the β_2_ adrenergic receptor (GPCRmd ID: <u>121</u>, PDB ID: 4LDE) as an example and focusing on Y326^7x53^ from the NPxxY motif to illustrate the inactivation process from an active structure. The figure displays three visualization modes for this residue: the first one shows the variation over time of the Cα of Y326^7x53^, reflecting three distinct states: active, intermediate, and inactive, where a complete flip of the Y326^7x53^ side chain orientation is seen because of the inactivation. The second mode, named one-dimensional CS, displays the mean predicted signal of the side chain atoms of Y326^7x53^ as a 1D-NMR spectrum. The two-dimensional CS shows the distribution of CSs for the Cα and Hα of Y326^7x53^, mimicking a 2D-NMR spectrum, where the predicted CS signal illustrates the different states of the (in)activation process of the GPCR.

Built on our GPCRmd database, a large-scale collection of high-quality and unified GPCR MD data, we have computed the chemical shift evolution over 970 GPCR MD systems belonging to 400 experimentally solved structures of GPCRs in different functional states (e.g., ligand-bound complexes and apo receptors, see **Figure 1B**). This dataset illuminates the chemical shift fluctuations along with the structural dynamics at a 0.2 ns resolution. In total, our chemical shift exploration of over 2.3 milliseconds of accumulative MD simulation time has yielded 7.2 million conformational snapshots with assigned chemical shift profiles and 28 billion CS values (**Figure 1B**). This data is made freely accessible via GPCRmd-NMR (https://www.gpcrmd.org/gpcrmd-nmr-tool) which is equipped with tailored web-based streaming, visualization, and data analysis tools (**Figure 1C**).

### GPCRmd-NMR aligns with experimental NMR observations

We have challenged the real-life performance of GPCRmd-NMR in the context of solid-state and solution NMR spectroscopy. For this, we have picked up two GPCR systems for which high-quality NMR data were available. First, we have validated the performance of our approach on the human bradykinin 1 receptor (B1R)^23^, a peptide receptor that plays crucial roles in chronic pain and inflammation. In the first step, we performed MD simulations of the B1R-peptide agonist complex (3 replicates of 500 ns). During MD simulations, the proline 8 (P8) residue of the peptide adopts a fully embedded configuration as confined by the peptide binding cavity during the MD simulation (GPCRmd ID: <u>2095</u>, PDB ID: 7EIB). We then reconstructed using GPCRmd-NMR the 2D ^13^C-^13^C double-quantum single-quantum (DQ-SQ) spectral pattern for all carbon atoms of the P8 residue of the peptide ligand (**Figure 2A**). Previously, the chemical shift data of the peptide agonist DAKD in complex with B1R were obtained via DNP-enhanced solid-state NMR spectroscopy^24^, which were collected at low temperature (ca. 100 K) and therefore present the conformational distribution of the bound ligand. We find that both the predicted (multi-colored) and experimental (dark blue) NMR signals of P8 overlay rather well (**Figure 2A, right**), indicating that our method aligns the structure plasticity with NMR observations. Additional 500-nanosecond replicate simulations reproduce an identical 2D ^13^C-^13^C DQ-SQ spectrum (**Figure S1**). The success of reproducing a small-sized low-temperature NMR dataset encouraged us to explore more comprehensive NMR data streaming at more physiological temperatures.

**Figure 2.**
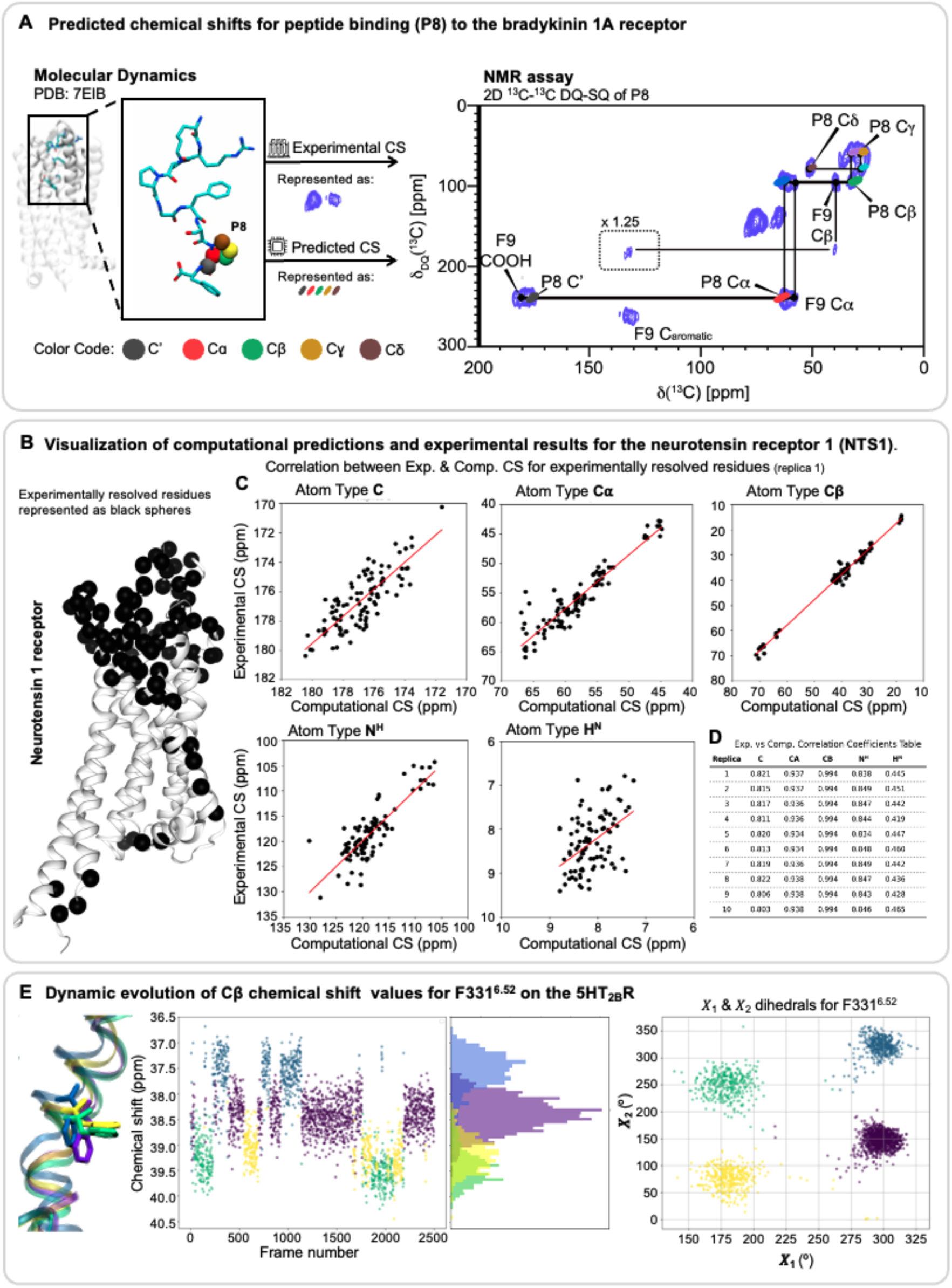
Benchmarking computational predictions against experimental NMR Chemical Shifts. **(A)** Predicted chemical shifts for peptide binding (P8) to the bradykinin 1A receptor (GPCRmd ID: <u>2095</u>, PDB: 7EIB). The signals from the experimental conditions are depicted as blue distributions in the 2D 13C-13C double-quantum single-quantum (DQ-SQ) spectrum. We overlapped the predicted CS signals for the C’ (grey), Cɑ (red), Cβ (green), Cɣ (orange), and Cδ (brown) atoms. We observe a notable overlap between the experimental and computational signals. (**B**) Visualization of computational predictions and experimental results for the neurotensin receptor 1 (NTS1). Amino acids of NTS1 where experimental NMR backbone chemical shift assignments could be obtained are shown as black spheres, most of them located in the extracellular region forming the orthosteric ligand binding pocket taken from Mohamadi *et al.^25^*(BMRB ID: 51907, GPCRmd ID: <u>2097</u>, PDB ID: 4BWB). (**C**) CS correlation plots for each atom type between computational (x-axis) and experimental (y-axis) CS values for replica 1 (see Figure S5-S9 for replica plots 1 to 10). We see a strong correlation for the heavy atoms, and a lower correlation for hydrogen atoms. (**D**) Table of correlation coefficients for all replicas 1 to 10 and atom types. The correlation coefficients are maintained between replicas, suggesting the robustness of combining MD simulations with CS predictions. (**E**) Dynamic evolution of Cβ chemical shift values for F331^6.52^ on the 5HT_2B_R (GPCRmd ID: <u>85</u>, PDB ID: 4IAQ). This residue exhibits four distinct conformational states over the MD, differentiated by the 𝛸_1_ and 𝛸_2_ dihedral angles. We used the same color-coded clustering scheme to visualize the clusters in the CS evolution over time.

We then moved to a case of solution NMR spectroscopic study on *rattus norvegicus* (rat) neurotensin receptor 1 (NTSR1). The NTSR1 receptor senses and responds to the neuropeptide neurotensin, which modulates dopamine signaling, regulates body temperature, and influences pain perception. Mohamadi and coworkers^25^ performed extensive backbone chemical shift assignment on this receptor bound to the natural agonist neurotensin-1. The dataset contains CS information on backbone signals, including the atom types C’, Cɑ, Cβ, N^H^, and H^N^ (BMRB ID: 51907). We used this rich dataset to evaluate systematically the performance of CS predictors. We conducted MD simulations (10 replicas of 1 μs, GPCRmd ID: <u>2097</u>, PDB ID: 4BWB) mirroring the experimental conditions used for recording the NMR data^25^.

Our results illustrate an excellent agreement between computational predictions and experimentally determined chemical shifts for the targeted residues (**Figure 2B-C**), with strong correlations observed for heavy atoms (R^2^ = 0.80 to 0.99) (**Figure 2D**). In contrast, predictions for H^N^ atoms show lower correlations (R² = 0.42 to 0.46), which is expected given that ^1^H chemical shifts are more sensitive to subtle inaccuracies in structural or electronic modeling^26^. Notably, the consistent correlation across replicates further underscores the robustness of integrating chemical shift predictors with molecular dynamics simulations (**Figures S2–S6**).

Our finding emphasizes the value of computational methods, particularly in scenarios where conventional NMR experiments cannot provide sufficient information for a reliable resonance assignment. By leveraging the consistency between computational and experimental data, these methods emerge as a promising tool for resonance assignment in regions that may be elusive for traditional NMR approaches due to the large size of the protein-membrane system.

### Illuminating underexplored GPCR dynamics via GPCRmd chemical shift streaming

Beyond conventional averaged biophysical parameters or their distributions from spectroscopic studies, our advanced computational framework enables the continuous tracking of spectroscopic parameters throughout the full-atomistic structural evolution (**Figure 1C**, upper panel). This approach is particularly valuable in challenging cases where chemical shift values arise from highly heterogeneous structural ensembles. Here, we demonstrate this capability for the 5HT_2B_R - a promising drug target for pulmonary arterial hypertension, valvular heart disease, and related cardiopathies^27^. Monitoring the Cβ chemical shift evolution of the F311^6x52^ (**Figure 2E**), a functionally relevant residue in the ligand binding pocket, we find that it samples four distinct conformational states in the apo 5HT_2B_R over time (GPCRmd ID: <u>85</u>, PDB: 4IAQ^28^). The conformational exchange over these four microstates occurs at the nanosecond scale and would typically result in one averaged signal when studied by NMR spectroscopy. Such a case suggests that a computational framework like GPCRmd-NMR can reveal underexplored structural dynamics by integrating MD simulations with the computation of relevant biophysical readouts. This is demonstrated in a real case in the following.

Taking the example of NTS1R as studied by Bumbak and co-workers using solution NMR spectroscopy^29^, we have reconstructed the 2D ^1^H-^13^C NMR spectroscopy of methionine methyl groups in this receptor via our GPCRmd-NMR approach. To explore how changes in receptor structure induce chemical shift alterations in the NTS1R, we performed MD simulations (3 replicas of 1 μs) for apo-NTS1 (GPCRmd ID: <u>2099</u>, PDB ID: 6Z66) and computed the chemical shift profile. Focusing on the methionine residues in the transmembrane (TM) helices TM4 (M204^4x61^), TM5 (M244^5x46^, M250^5x51^) and TM6 (M330^6x57^), the NMR spectral patterns of their methyl groups synthesized by GPCRmd-NMR resemble the experimental observations (**Figure S7**). A clear right-shift of M250^5x51^ compared to M330^6x57^ can be well-reproduced by GPCRmd-NMR (**Figure 3A-B)**, indicating that our MD simulations retain the structural differences of these two sites. Intriguingly, GPCRmd-NMR revealed that the observed average chemical shift (∼1.70 ppm) of M250^5x51^ methyl group arises from two distinct conformations (**Figure 3C**), in which this methyl group orients towards either the membrane or the receptor interior. The average of the predicted chemical shifts (δ(1H) ∼1.70 ppm) (**Figure 3B**) matches well the experimental value (δ(1H) ∼1.68 ppm) (**Figure 3A**). This finding uncovers a previously unrecognized rapid conformational switching near key receptor activation sites, highlighting dynamic features that are critical for receptor function yet elusive to conventional experimental approaches. Notably, the rich structural and spectroscopic insights offered by GPCRmd-NMR across the 3D GPCRome can be harnessed to explore undercharacterized GPCR dynamics in a highly flexible and customizable framework.

**Figure 3.**
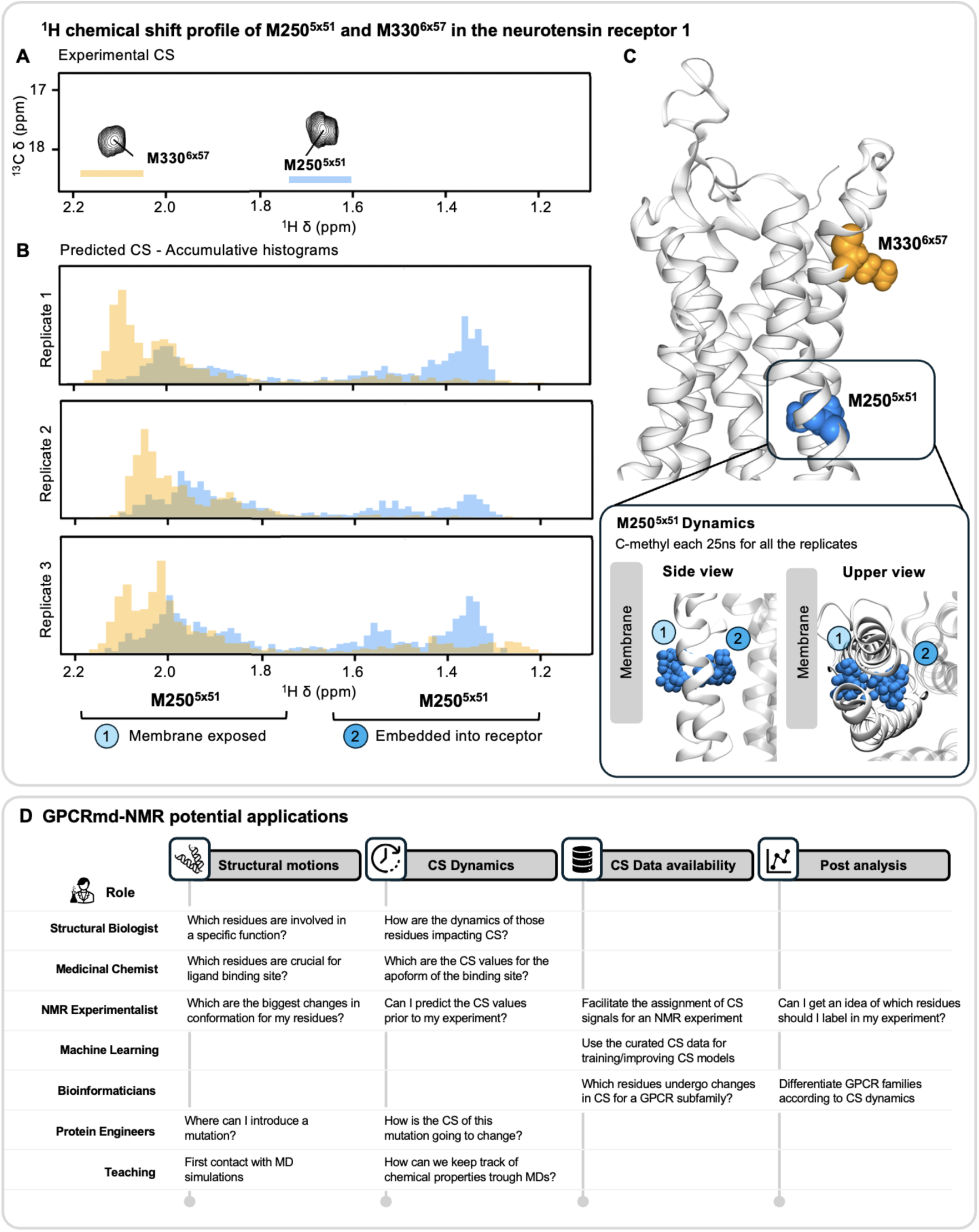
Linking conformational states and chemical shift variability in NTS1 methionines and GPCRmd-NMR potential applications. **(A)** ^1^H chemical shift profile of M250^5x51^ and M330^6x57^ in the neurotensin receptor 1. Experimental CS data for M250^5x51^ and M330^6x57^ extracted from Bumbak et al. ^29^, Figure 2). (**B**) Predicted CS data for M250^5x51^ and M330^6x57^ based on 3 x 1 s simulations. The predicted CS is set to match the axis of the experimental CS, and we plot the cumulative histogram of the CS values for the two methionines. Notably, M330^6x57^ exhibits a single distribution in the histogram, while M250^5x51^ manifests two distributions. (**C**) Structural depiction of the neurotensin receptor 1 and the location of M250^5x51^ with two different side chain conformations (blue spheres) and M330^6x57^ (orange spheres). (**D**) Potential applications for the Chemical Shift tool for different backgrounds.

GPCRmd-NMR also permits resolving such dynamic conformational plasticity on higher spectroscopic dimensions. Our portal is highly flexible for constructing any of such spectroscopic representations. The most studied atoms in NMR protocols are available as pre-defined selections, including methionines (^13^Cε^1^Hε), isoleucines (^13^Cδ^1^Hδ), alanine (^13^Cβ^1^Hβ), leucine (^13^Cδ^1^Hδ) and valine (^13^Cγ^1^Hγ). Ultimately, the 2D chemical shift plots are interactive, allowing users to select specific residues directly within the plot. Upon selection, the chosen residues are highlighted and zoomed in the GPCRmd viewer. In the streaming mode, users can track the simulation frame as it progresses, facilitating the correlation of chemical shifts with specific frames of the MD simulation. In **Figures S8-S12**, we provide an overview of the chemical shift tool and the streaming chemical shift mode.

### Integrative enhancement of spectroscopic pipelines via GPCRmd-NMR

Though being pivotal to accessing GPCR dynamics, the current spectroscopic approaches to GPCRs suffer from several key bottlenecks spanning over their full pipeline, namely the (i) signal assignment, (ii) reporter selection and (iii) integrated hypothesis-generation and spectroscopic solutions. Encouraged by the abovementioned performance of GPCRmd-NMR, we have envisioned several new computation-driven approaches that could immediately boost these spectroscopic efforts on GPCRs.

An important strength of GPCRmd-NMR is to resolve the inherent spectral complexity of GPCRs dynamics and by this significantly simplifies the process of NMR signal assignment. Here we provide a user-friendly solution based on the developed chemical shift predictor framework. Aiming to facilitate this task, we mimic the traditional 1D and 2D NMR spectra plots, which allow users to examine and differentiate signals within the same plot. To illustrate their utility, we focus on the β_2_ adrenergic receptor - a GPCR that plays a critical role in managing asthma and respiratory disorders through bronchodilation, mucociliary clearance, and anti-inflammatory effects^30^. Specifically, we examine the Cα and Nitrogen atoms of the backbone of two distinct aspartates (**Figure 4A**). One is situated in the helical region of the transmembrane domain TM2 at the functionally highly relevant allosteric sodium binding site (D79^2x50^) and the other in the extracellular loop 2 (D192^45x51^) - a loop known to be involved in ligand binding in the β_2_ adrenergic receptor^31^. The differences in location and chemical environment between these aspartate residues is reflected in different chemical shift values (**Figure 4B-C**). For a more detailed explanation of these visualization modes, including their significance and interpretation, refer to **Supplementary Note 1**.

**Figure 4.**
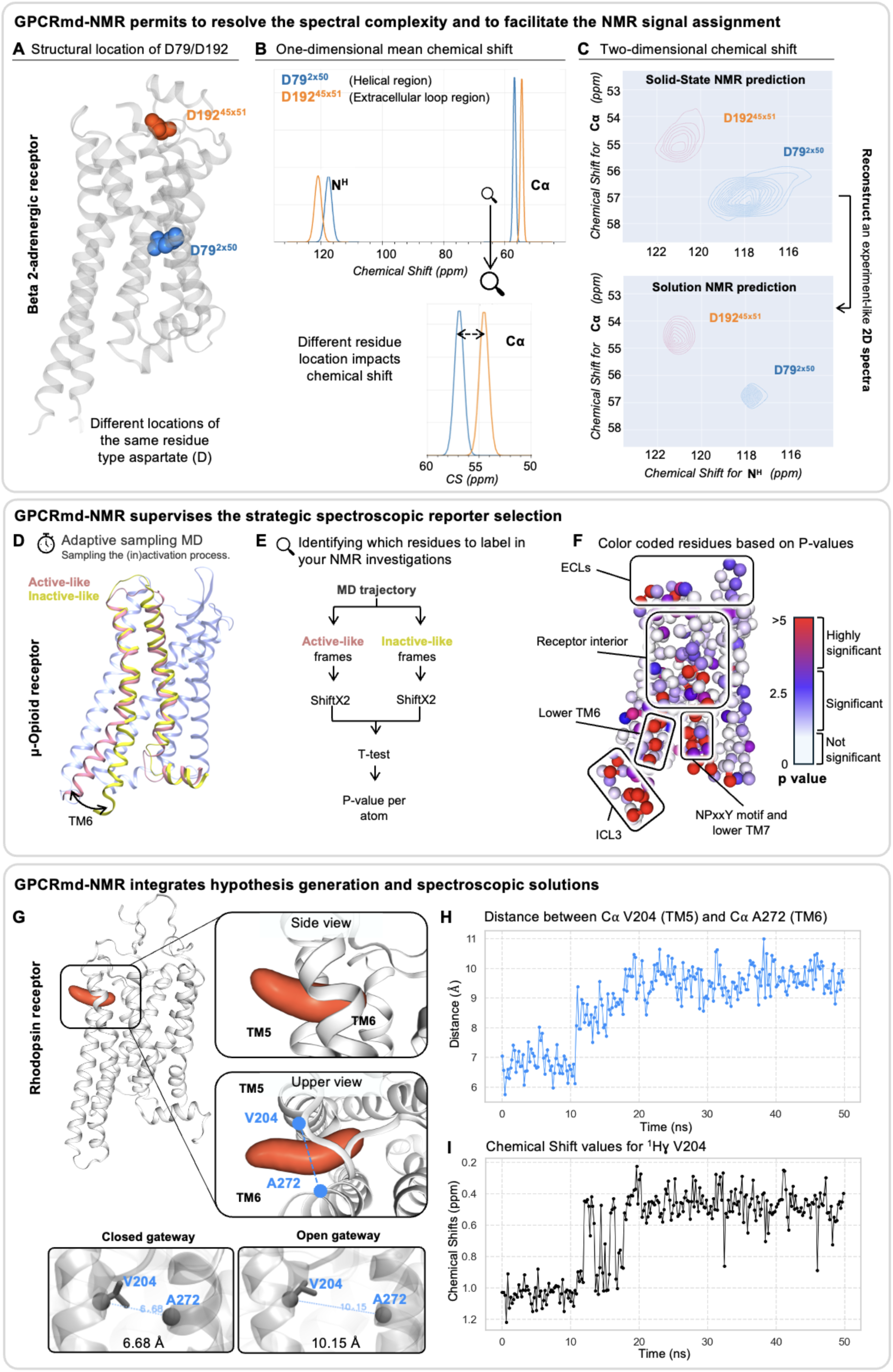
Integrative enhancement of spectroscopic pipelines via GPCRmd-NMR. (**A**) β_2_ adrenergic receptor with the two selected aspartates residues, D192^45x51^ located on the extracellular side, while D79^2x50^, a highly conserved residue, faces the middle of the receptor. (**B**) One-dimensional chemical shift for D79^2x50^ and D192^45x51^. The mean signals show no clear overlap, indicating that the two signals are significantly different. (**C**) The two-dimensional chemical shift mode includes two predictions: solid-state NMR, where the distributions of the computed chemical shift values are displayed, and solution NMR spectra, where the means of the distributions for both residues are displayed. (**D**) Classification of active-like (pink) and inactive-like (yellow) structures for the μOR based on the inwards-outwards movement of the TM6. (**E**) Schematic workflow of CS computation using SHIFTX2. (**F**) The μOR structure visualizes the color-encoded p-values for the CS values of the nitrogen atoms in the backbone of each residue. Red represents highly significant changes; blue indicates significant differences white denotes residues with no discernible changes in CS signals between the inactive and active μOR receptor. (F) Density map (orange) of the inserted lipid tail within the cryptic pocket of the Rhodopsin receptor (GPCRmd ID: <u>872</u>, PDB ID: 5DYS)**. (H)** Distance evolution between the C⍺ atoms of V204 and A272 (blue). (**I**) Evolution of the 1Hγ chemical shift for V204 (black), with a correlation value of 0.83 with the distance between V204 and A272.

Furthermore, GPCRmd-NMR supervises the strategic spectroscopic reporter selection. In any spectroscopic studies of protein dynamics, the selection of suitable reporters is of crucial importance. However, this is often done via human experiences obtained via the costly try-and-error processes. GPCRmd-NMR could explore a large set of spectroscopic reporters and provide a systematic evaluation over the inexhaustible options in the real-life setting. Taking the μ-opioid receptor (μOR), a target for most clinically and recreationally used opioids^32^, as an example (GPCRmd ID <u>199</u>, PDB ID: 5C1M, submitted to GPCRmd by Kapoor et al.^33^), we aimed to identify reporter positions with significant readouts for studying receptor (in)activation. In a first step, MD frames were categorized into active and inactive states based on the receptor conformation (see method section, **Figure 4D**) and backbone ^15^N chemical shift values were obtained for both states (**Figure 4E**). As expected, we observe large chemical shift changes occur mainly in regions of known functional relevance (**Figure 4F**). For example, alterations occur around the NPxxY motif in the lower TM7 (L331^7x48^, L335^7x52^, Y336^7x53^) (blue to red) due to (in)activation motions. This is also evident for residues in TM6 with pronounced differences in V285^6x40^ and I278^6x33^ or the intracellular loop 3 (ICL3) because of the TM6 in/outward movement. Most importantly, GPCRmd-NMR also reveals additional regions within the receptor interior that exhibit substantial backbone chemical shift changes (e.g., Y326^7x42^), which are not apparent in static structures and may point to previously unexplored functionally relevant receptor areas. Our approach can be extended feasibly to other receptors, functional states, sites and spectroscopic reporters, therefore paving the way for rational design of biophysical approaches for studying GPCR dynamics.

Finally, GPCRmd-NMR integrates hypothesis generation and spectroscopic solutions. Recent computational studies have demonstrated that we can monitor the behavior of entrance gateways with MD simulations in GPCRs^16^. These transient pockets form and disappear dynamically. In this case study, we explore the formation of entrance gateways by using predicted CS values on the rhodopsin receptor, a key receptor for dim-light vision ^34^. To this end, we analyze an MD simulation (GPCRmd ID: <u>872</u>, PDB ID: 5DYS) where we observe the spontaneous insertion of a membrane lipid’s aliphatic tail into the receptor, specifically between the upper regions of transmembrane helices TM5 and TM6. This event occurs during the first 50 ns of the simulation, capturing the transition from an unbound lipid state to the moment when the lipid tail fully penetrates the receptor.

We evaluated the correlation between the predicted chemical shifts (CS) of the residues near the entrance gateway and the distance between the C-alpha atoms of residues V204 and A272, defining the opening of the cavity (**Figure 4G**). As observed in **Figure 4H-I**, V204 displayed a strong correlation (R² = 0.83) for its 1Hγ chemical shift, an atom that can be easily labeled experimentally. The 1Hγ chemical shift evolved from approximately 1.2 ppm in the initial simulation steps (cavity closed) to 0.4 ppm once the lipid was fully inserted. While the gateways frequently open and close, this finding could be a starting point for exploring potential applications of NMR experiments to detect and localize the formation of entrance gateways in protein structures.

## Discussion

The decoding of complex protein dynamics is a long-lasting challenge in protein science. This task is particularly relevant for understanding the molecular mechanisms of GPCRs, as the cellular behaviors of these important drug targets depend strongly on the conformational space sampled by their distinct functional states. To tackle such a challenge, we have developed a new approach integrating *in silico* simulations and high-resolution experimental biophysical data at the 3D GPCRome level. Built on a community-driven, open-access GPCRmd platform and showcased on several receptors, our toolkit provides new insights into GPCR dynamics and enhances their study from multiple perspectives.

First, GPCRmd-NMR improves the biophysical data interpretability and functional state assignment. A major practical hurdle in the NMR studies of protein dynamics is the data interpretation. Chemical shifts, though highly sensitive to protein structure and environment, are difficult to directly translate into structural parameters. Many pioneering works have targeted this challenge, yet it becomes pronounced in highly dynamic proteins such as GPCRs^4^. GPCRmd-NMR enables the dynamic streamlining of chemical shift evolutions at high temporal resolution. As showcased on hB1R-peptide (**Figure 2A**), GPCRmd-NMR illuminates the complex conformational distribution together with the NMR data on a frozen sample. On NTS1R at a more physiologically relevant temperature, our approach permits us to unravel the chemical shift distribution averaged by the dynamic sampling of protein conformations. (**Figure 3A-C**). Such a capacity in interpreting the static and averaged NMR data based on the time evolution of protein structures will open the way for uncovering the functional states encoding these biophysical readouts. Despite the success of many NMR reporters (e.g. ^19^F-containing groups, ^13^C-methyl groups, backbone ^1^H-^15^N amide groups) in GPCR studies, they suffer from the limited information content (monoplexing approach) and/or the lack of correlation information between different sites (even in multiplexing methods). GPCRmd-NMR comprehends these valuable experimental biophysical data by “projecting” them back to the full atomistic protein dynamics.

Second, GPCRmd-NMR illuminates the underexplored GPCR dynamics via the MD “mining” of NMR data. As discussed above, the experimental biophysical data capture only some local or low-dimensional structural features. As demonstrated in the hB1R-peptide case (**Figure 2A**), our MD interpretation of the NMR readouts has revealed complex structural plasticity that cannot be derived feasibly by conventional static NMR data analysis approaches. This capacity to illuminate the hidden structural dynamics has become even more evident on NTS1R, where a local sidechain conformational switch located next to the receptor activation site can be unmasked by the GPCRmd-NMR analysis of the experimental chemical shift values (**Figure 3A-C**). Our GPCRmd-NMR toolkit provides new opportunities for identifying the previously unnoticed function-relevant sites by mining the experimental biophysical data.

Third, our GPCRmd-NMR platform accelerates the generation of hypothesis-driven biophysical strategies for mapping function-relevant receptor dynamics. A prerequisite for the success of biophysical studies of receptor dynamics is the proper selection of the types and the specific locations of reporters. Traditionally, these biophysical reporters have been chosen based on prior experience or through laborious try-and-error processes, presenting a major bottleneck of the biophysical studies of receptor dynamics. As shown on rhodopsin, GPCRmd-NMR can generate function-relevant structural hypotheses (e.g., transient cryptic channels involved in ligand entry) and bootstrap potential spectroscopic solutions by exploring the structural evolution of receptors. This feature not only significantly reduces the cost and risk of the biophysical studies but also guides the design of new biophysical strategies to probe previously underexplored functional sites. Additionally, GPCRmd-NMR establishes *in silico* biophysical profiles across the 3D GPCRome, providing baseline patterns that facilitate the selection of receptor-specific biophysical reporters that map the unique structural features of each receptor.

Lastly, GPCRmd-NMR presents a template for the open-access integration of MD simulations and diverse experimental biophysical data. As discussed, GPCRmd-NMR boosts the NMR studies of GPCR dynamics across the entire research pipeline, from hypothesis generation and reporter selection to data interpretation. Its open database architecture permits incorporation of emerging deep learning-based CS predictors and quantum-chemical level chemical shift calculations. Together with the more vigorous sampling of conformational space, they could resolve some remaining limits raised by the currently implemented chemical shift predictors, for example, the relatively poor ^1^H chemical shift performance. Thanks to the high spatiotemporal resolution of full-atomistic MD at the backend of our platform, the demonstrated success on supporting NMR spectroscopy can be expanded to many other spectroscopic methods (e.g. DEER, smFRET, UV-Vis, fluorescence), biophysical parameters (e.g. distances, wavelength, signal intensity) or even biochemical readouts (e.g. H/D exchange). Some biophysical data focusing on slower millisecond-scale dynamics, typically beyond the reach of common MD simulations, can be resolved by combining MD data of the same receptor for multiple states, an advantage of GPCRmd. Our platform provides biophysical method developers new opportunities for side-by-side benchmarking using real-life GPCR examples. The integrative power of GPCRmd will also catalyze the joint development of complementary experimental methods and biophysical predictors/calculation tools via a shared MD dataset. GPCRmd-NMR is specifically tailored to meet the needs of scientists from diverse disciplines, such as medicinal chemists and NMR-based structural biologists, while also providing valuable resources for ML engineers and bioinformaticians. Altogether, this broad accessibility makes the GPCRmd CS tool a valuable resource for advancing research across multiple directions in GPCR research (**Figure 3D**). Moreover, this framework can serve as a template for similar efforts in other important protein families such as kinases, chaperones, nuclear receptors, ion channels and transporters.

### Data availability and web server access

CS widget can be accessed online in the web https://www.gpcrmd.org/gpcrmd-nmr-tool by selecting any of the systems available.

### Open Access Scripts for Community Use

All scripts employed in our analysis are freely available to the community, providing a foundational codebase for researchers to reproduce experiments following our presented guidelines (see methods “Reproducibility and Data Sharing”).

## Materials and Methods

### Data processing for CS computation

The computing protocol is composed of an initial data preprocessing part, where non-protein atoms and non-standard amino acids are excluded from the prediction. Next, CS values are computed by a python pipeline and the binaries of the programs ShiftX2 (version 1.10) and Sparta+ (version 2.90). The pH and temperature parameters used by the predictors were set to 7 and 300 kelvins, respectively. This was done to be in line with the MD parameters used to run the simulations in GPCRmd. Afterwards, the statistical error from the prediction is computed. For this, we calculated a block average analysis for each of the atoms included in the prediction. This blocking strategy allows us to treat data obtained from MD simulations as a sum of uncorrelated random variables, obtaining an accurate variance estimate for each predicted CS value. To ease up the data comprehension, we provide the error as the sum of squares of the previously mentioned block average analysis and the intrinsic error of the predictor, defined by the authors^21,22^. For the 2D solution NMR plot, we represented the CS distributions by calculating the mean and deviation from the original 2D NMR data, aiming to mimic an average distribution of experimental solution NMR. Lastly, the data is rearranged into a user-friendly format ready to be downloaded from the server. This process usually takes around 5-8 hours of computing per system using our facilities and the code is scalable to any future server upgrades.

### Data visualization

The resulting CS predictions are stored in CSV format, and they are used to produce the plots in the three different visualization modes. CSV format is easy to read and manipulate with most of the analysis tools available in the market. Two distinct CSV files are available for download from the server: a raw dataset and a simplified dataset. The raw dataset contains predicted CS values for each frame of the trajectory, while the simplified dataset includes the mean values and associated errors of these predictions. This difference was meant to avoid downloading unnecessarily large datasets. Each of the lines in both datasets represents a single atom of the system, including the information of its atom and residue ID.

The interactive plots were generated by using the python libraries *plotly* (https://plotly.com/) and *bokeh* (https://bokeh.org/). By selecting any of the atoms shown in the plots, a zoomed-in representation will appear in the GPCRmd viewer.

### Molecular Dynamic simulations

CS values were computed for three specific case studies analyzed in this study as well as for the entire GPCRmd repository.

#### 1. Preparation of MD simulations for case studies

*B1R study:* For the B1R, modeling was not necessary as the structure was directly obtained from the Protein Data Bank (PDB ID: 7EIR), whereas for the NTS1, we used two different models, adapting each to the structure and models used in their corresponding NMR experiments.

*NST1 for Bumbak et al. study.* For the work conducted by Bumbak *et al.*, the initial structure of the NTS1 was based on PDB ID 6Z66, spanning from Y62 to N370. Using Prime from Maestro, we modeled the intracellular loop 1 (ICL1, K92 to S97) and extracellular loop 2 (ECL2, S214 to G222). A homology model was subsequently generated using the target sequence outlined in the supplementary material of the authors’ paper, incorporating their key mutations (M181L, M267L, M293L, and M408V). An additional step of minimization of the loops was performed using Prime, with simultaneous minimization of protein hydrogen bonds to accommodate the modeled residues and prevent clashes.

*NST1 for* Mohamadi *et al. study:* For the NTS1 case study we used PDB ID 4BWB, and no additional modeling was required.

##### General system setup

All system setups were conducted using the CHARMM-GUI server. Protein cappings were achieved using methyl groups to neutralize them (ACE & CT3), while peptide cappings were intentionally left uncapped. Protonation states of amino acids were determined using PropKa 3.0 at a pH of 7. For the NTS1, a disulfide bridge was formed between C225 and C142. The receptors were embedded into a homogeneous bilayer composed of Palmitoyl-oleoyl-phosphatidylcholine (POPC) after aligning the system with the OPM server. The systems were neutralized and ionized by adding appropriate concentrations of NaCl and MgCl2 to mimic experimental conditions. For the B1R, we used salt concentrations of 0.10 M NaCl and 0.05 M MgCl2, and for the NTS1, we used 0.15 M NaCl.

The software used for MD simulations was ACEMD3.3 using the CHARMM36m all-atom forcefield. Each system was subjected to energy minimization for 500 steps to alleviate steric clashes and optimize atomic positions. This was followed by a 20 ns equilibration phase where the system temperature was gradually increased from 0 to 300 kelvin and then equilibrated in an NPT ensemble. To stabilize the system, restraints were applied to C-alpha atoms, gradually released from 1 kcal/mol to 0 kcal/mol over 15 ns, while restraints on remaining heavy atoms were relaxed from 0.1 kcal/mol to 0 kcal/mol. The production runs were performed using a timestep of 4 femtoseconds under the NVT ensemble at a constant temperature of 300 kelvins without restraints. The duration of the production runs varied based on the experiment conducted:

● For the B1R study, three replicates of 500 ns were performed.
● For the NTS1 study conducted by Mohamadi *et al*.^25^, ten replicates of 1 µs were performed.
● For the NTS1 study conducted by Bumbak *et al*., three replicates of 1 µs were performed.

#### 2. MD simulations from GPCRmd

The computational details of the MD simulations at GPCRmd are described at: https://github.com/GPCRmd/simulation_pipeline. In addition, the users can review the specific details for each simulation on the server in the simulation report section.

##### Analysis and Visualization

For CS analysis, simulations were processed using the ShiftX2 and Sparta+ software to predict CSs for each trajectory. The specific pipeline used to compute and analyze the results for each of the experiments can be found in the Reproducibility and Data Sharing section. Below is a summary of the content for each section:

*1. Predicting CSs for Peptide Binding to the Bradykinin 1A Receptor:* Read the file from experimental DQ-SQ NMR and convert these values into a grid. Read the predicted CS files, transform the values to DQ-SQ format and overlap the 2 spectra into a single plot, coloring each atom type differently. Black lines were added to identify correlated pairs of atoms.
*2. Unveiling the Robustness and Potential of CS Predictors in Neurotensin Receptor Studies:* The experimental data used in this study is sourced from the BMRB databank. We calculate the correlation between the experimental CS and the average value of the predicted CS obtained from the MD simulations for each replicate. This process is repeated for every atom type, and the findings are presented in a tabular format. Additionally, we illustrate, using NGL representation^35,36^, the residues that contain experimental values.
*3. Going Beyond Static NMR Interpretation Considering Receptor Flexibility:* In this study, we delve into receptor flexibility by analyzing predicted CSs. We construct a histogram comparing the CS of the HE atom from the methyl groups of M250 and M330. Each distribution is depicted with a distinct color.
*4. Guiding Experimental Studies - Receptor (In)Activation of the μ-Opioid Receptor:* In this study, we analyzed a community-contributed 1.2 µs adaptive sampling simulation capturing the (in)activation dynamics of the µ-opioid receptor while bound to the agonist morphine (GPCRmd ID <u>199</u>, PDB ID: 5C1M, submitted to GPCRmd by Kapoor et al.^33^). This simulation allows us to explore the dynamic behavior and structural changes of this receptor during the activation process. To begin our analysis, we first read and align the MD trajectory frames uploaded in GPCRmd. We established as reference structures representing both the inactive (PDB ID: 4DKL) and active states (PDB ID: 5C1M) of the receptor. These references serve to categorize the frames with an RMSD of 1.5 Å or less from the reference structure into two groups: active and inactive-like. We perform a Welch’s t-test to determine the significance of differences in CS values per residue. Finally, we scale and map the p-values into B-factors within a PDB file for visualization purposes. The resulting data allows for the identification of the most variant residues between active and inactive forms, which can be readily visualized using software like VMD^37^, utilizing the B-factor as a color scheme. The complete code for this analysis is provided in a Jupyter Notebook file.

### Experimental NMR

The B1R was heterologously expressed and purified from Sf9-cells by using a baculovirus expression system. The B1Rwere solubilized in buffer (50 mM HEPES, pH 7.6, 150 mM NaCl, 5% (w/v) glycerol, 200 nM U–[^13^C,15N]–P8F9 DAKD supplemented with 1% n–dodecyl β–D– maltoside (DDM, Anatrace) and 0.1% cholesteryl hemisuccinate (CHS, Anatrace). The soluble B1R receptors were purified by using affinity purification Ni-NTA HisTrap HP column (GE Healthcare, Munich, Germany) with the help of Äkta system (GE Healthcare). The samples were eluted with buffer containing buffer (50 mM HEPES, pH 7.6, 150 mM NaCl, 5% (w/v) glycerol, 200 nM U–[13C,15N]–P8F9 DAKD supplemented with supplemented with 400 mM imidazole, 0.07% DDM, 0.007% CHS).

For DNP SSNMR sample preparation, the buffer of the purified protein sample was replaced with a buffer consisting of 50 mM HEPES–NaOD (pD 7.6), 150 mM NaCl, 5% (w/v) 13C–depleted and fully deuterated glycerol (Euriso–Top, Saint-Aubin, France), and 76% D2O (v/v). The buffer exchange was carried out using Amicon centrifuge concentrators (50 kDa cut-off, Merck Millipore, Darmstadt, Germany). During this process, U–[13C,15N]–P8F9 DAKD peptide was added in molar excess. The concentrated solutions were then supplemented with 10 mM AMUPol and mixed with 13C–depleted and fully deuterated glycerol in a 1:1 ratio. The mixtures were transferred to 3.2 mm sapphire or zirconium oxide SSNMR rotors using micropipettes and sealed with Vespel caps. The samples were subsequently frozen in situ in the low-temperature (ca. 110 K) gas flow inside the cryo-NMR probehead or directly in liquid nitrogen for storage.

All DNP-enhanced MAS ssNMR spectra were acquired using a Bruker Avance II 400 MHz (1H Larmor frequency) spectrometer, fitted with a 3.2 mm HCN triple-channel cryo-MAS probe head. The experiments were conducted at approximately 110 K under an 8 kHz MAS rate. High-power microwaves at 263.58 GHz were produced by a CPI gyrotron. The 15N–13C 2D TEDOR spectra were collected and processed using the same parameters as detailed in previous work . All spectra were externally referenced using a 13C-labeled alanine powder sample.

### Reproducibility and Data Sharing

All MD simulations used in this study, along with the necessary files to reproduce the experiments and input files, are available at GPCRmd. Users can visualize the simulations directly on the website and compute the CSs. Additionally, they can download the trajectories and the CS data. To facilitate the reproduction of the experiments, we uploaded the Jupyter-notebook files to the <u>GPCRmd GitHub repository</u> (https://github.com/GPCRmd/GPCR_Chemical_Shift/tree/main). These notebooks contain all the code, information and steps needed to replicate the results shown in this study. The user might need to download the data from the provided links in the Jupyter-notebook and place it into the proper directories, due to the size of the files.

## Supplementary Note 1

The first mode, termed “One-Dimensional Mean Chemical Shift”, presents a mean representation of the CS derived from the entire MD. The distribution’s center corresponds to the mean of the CS, while its width represents an error estimation for the CS prediction (refer to methods for more details). This error estimation guides the user to distinguish whether two measures are significantly different or not. In **Figure 4B**, a visual representation illustrates how the two distributions for the Cα and Nitrogen atoms between the aspartates differ in CS. For instance, the Nitrogen atom and the Cα of D79^2.50^ and D192^45x51^ are well separated in the β_2_ adrenergic receptor. The second mode, termed “Two-Dimensional Chemical Shift”, allows users to correlate two atom types by mapping their distribution on a plane, resembling a real 2D-NMR plot. This mode helps to quickly understand the ranges of CS values explored throughout the simulation. **Figure 4C** illustrates the “Two-Dimensional Chemical Shift” mode, where D79^2.50^ has slightly greater fluctuation in CS compared to D192^45x51^ for the Cα and Nitrogen atoms. Importantly, both visualization modes are interactive to enhance user comprehension. For instance, selecting a specific NMR signal allows users to visualize the corresponding atom within the 3D receptor structure using the GPCRmd viewer. A snapshot from GPCRmd highlighting the main features of both modes is presented in **Figures S8-S12**.

---

**Supplemental Figure 1.**
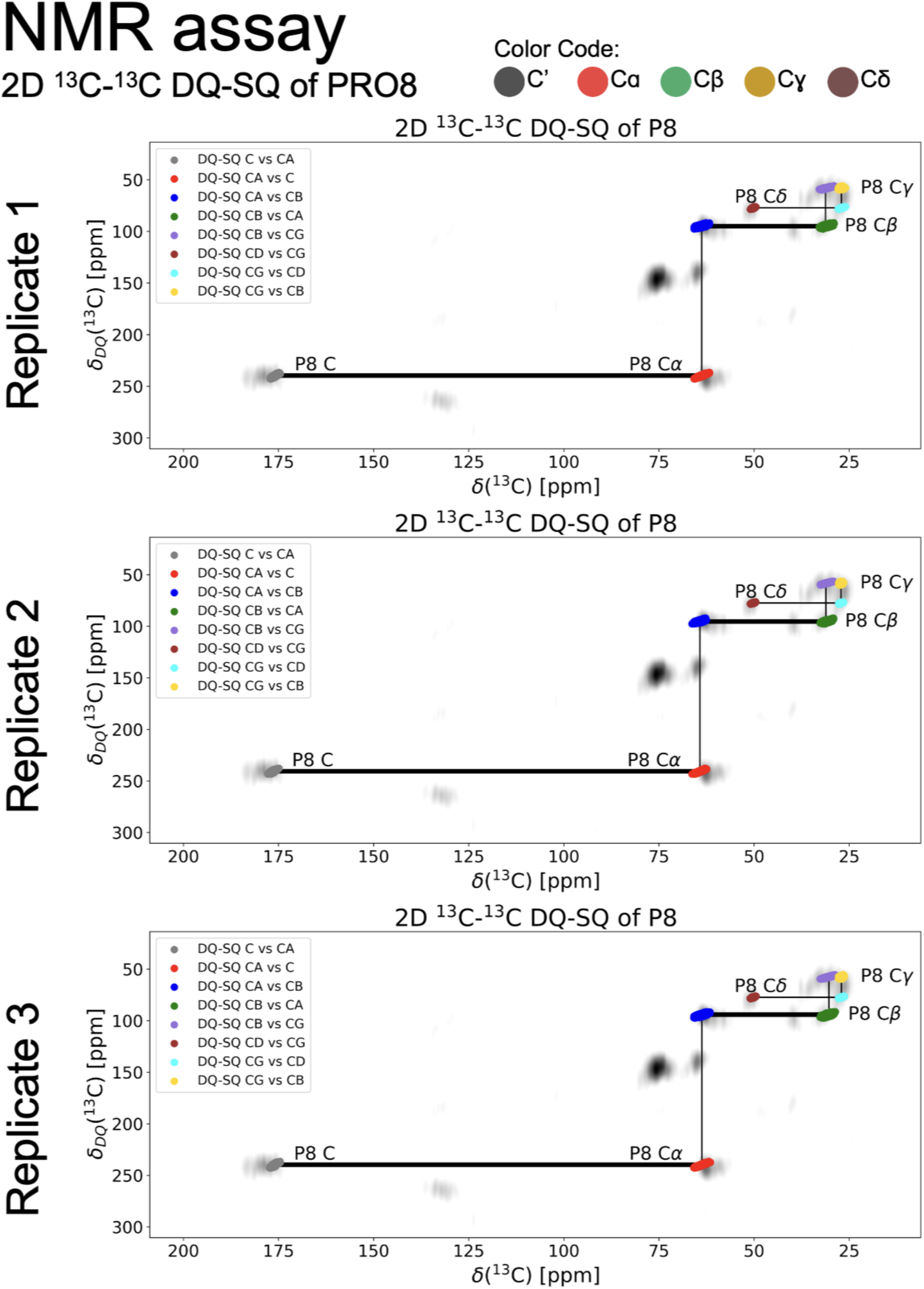
1^3^C-^13^C DQ-SQ overlap between predicted (colored) and experimental (grey) chemical shift values for P8 in the Bradykinin 1 Receptor. The lines between atom types represent the carbon-carbon correlations, indicating spatial proximity between them. Each replicate consists in a independent MD simulation of 500ns (GPCRmd ID: <u>2095</u>, PDB: 7EIB).

**Supplemental Figure 2.**
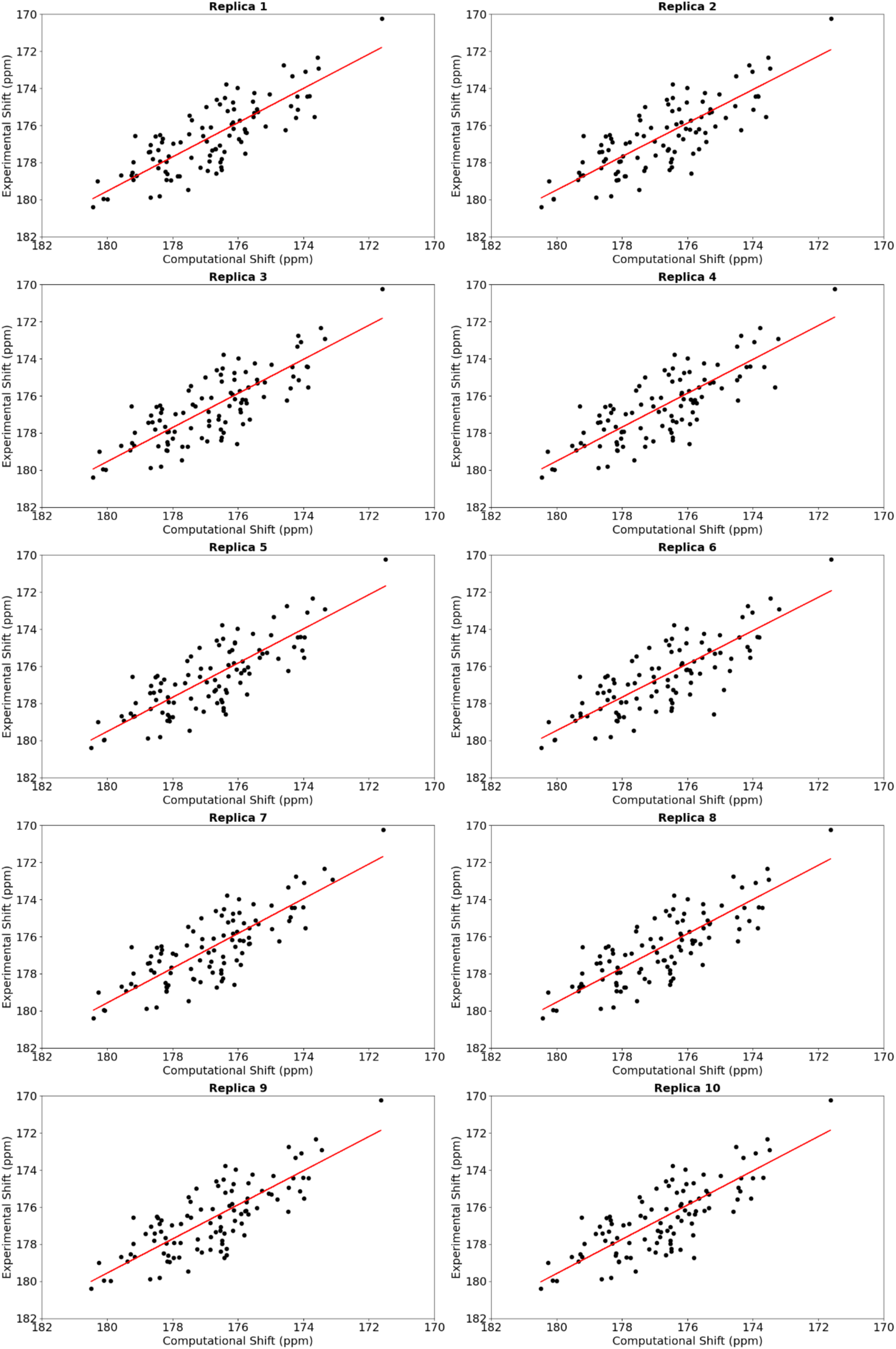
Chemical shift correlation for the C atom type between experimental and computational predictions for the neurotensin-1 receptor.

**Supplemental Figure 3.**
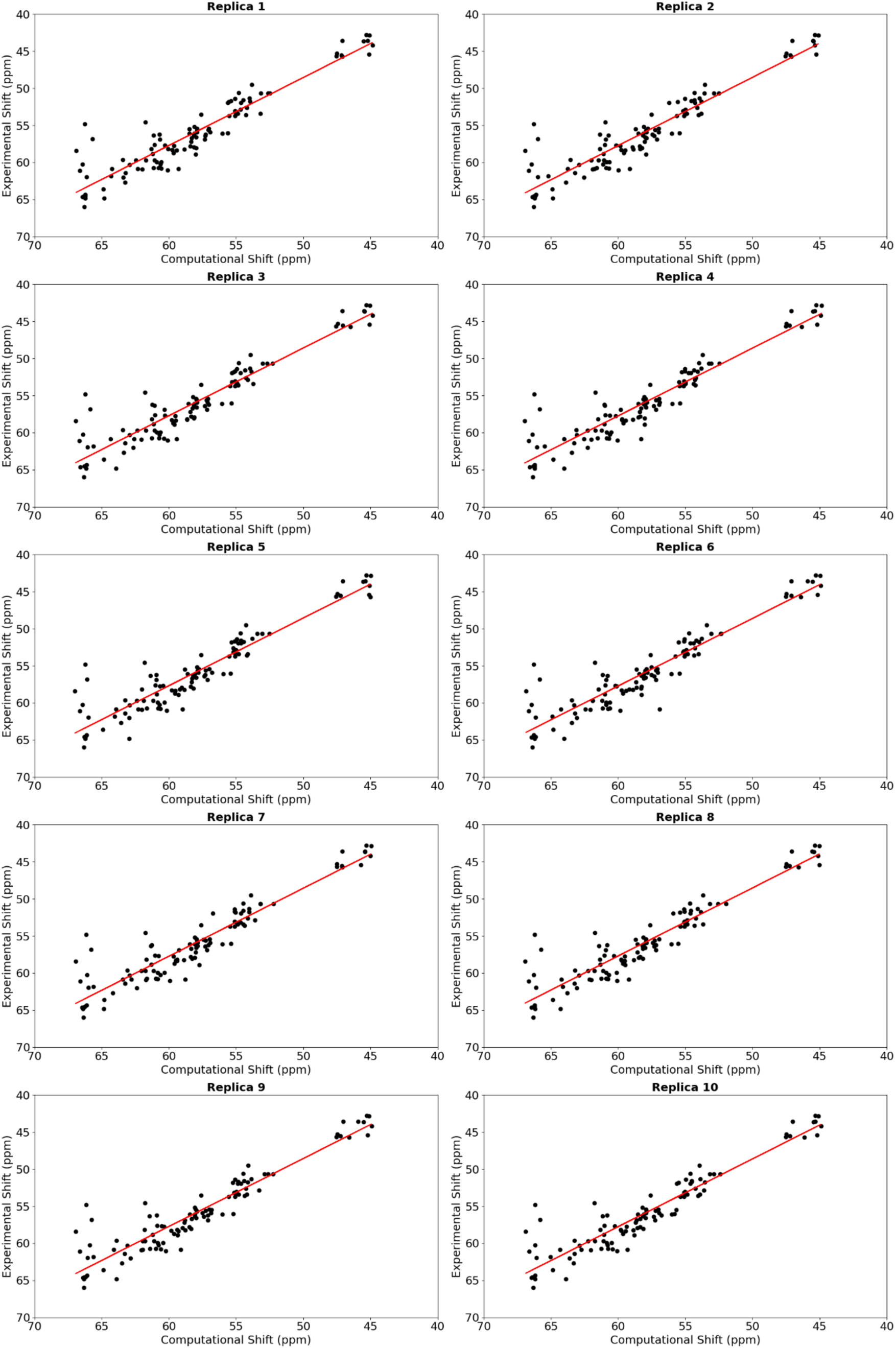
Chemical shift correlation for the Cɑ atom type between experimental and computational predictions for the neurotensin-1 receptor.

**Supplemental Figure 4.**
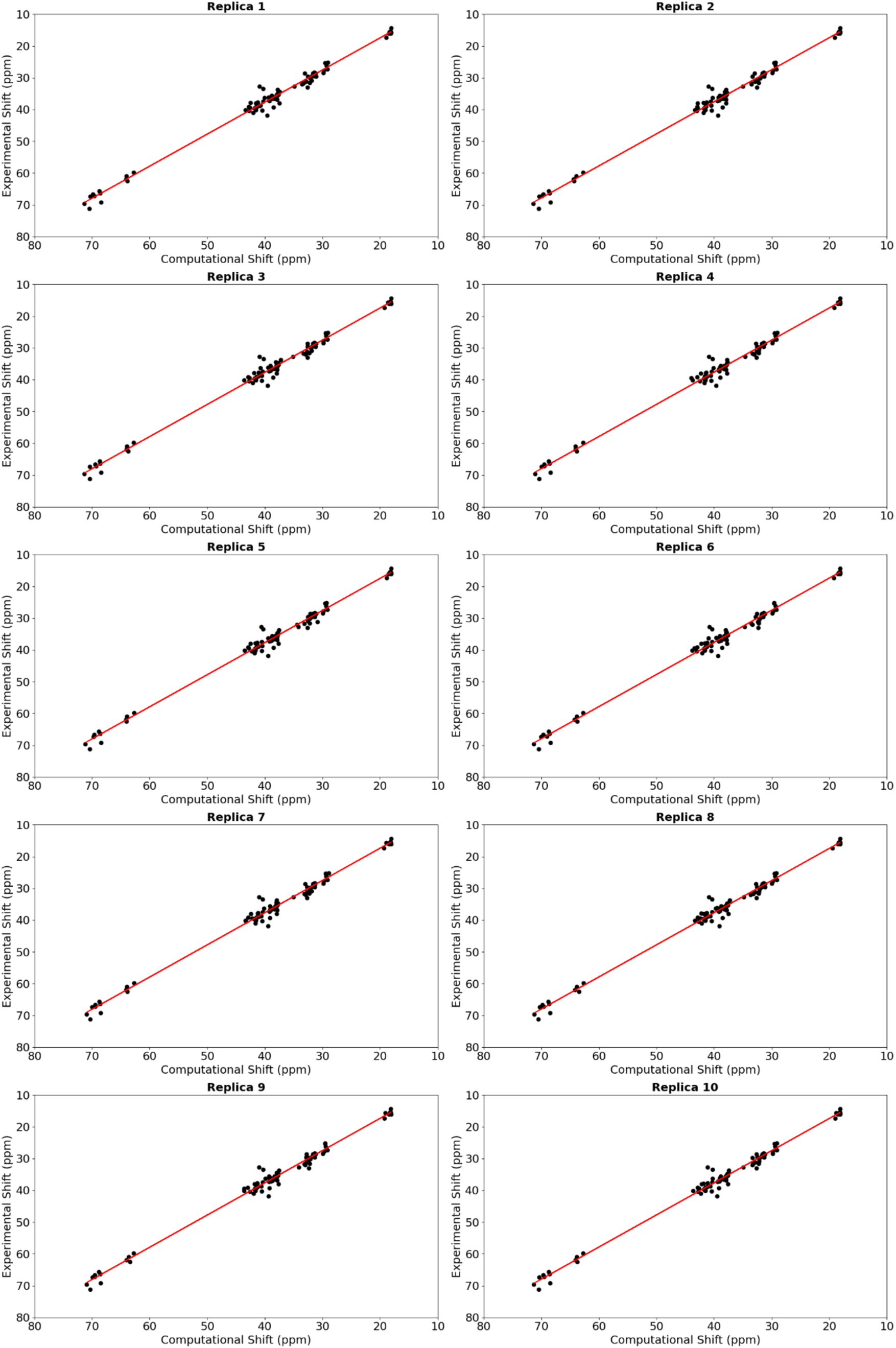
Chemical shift correlation for the Cβ atom type between experimental and computational predictions for the neurotensin-1 receptor.

**Supplemental Figure 5.**
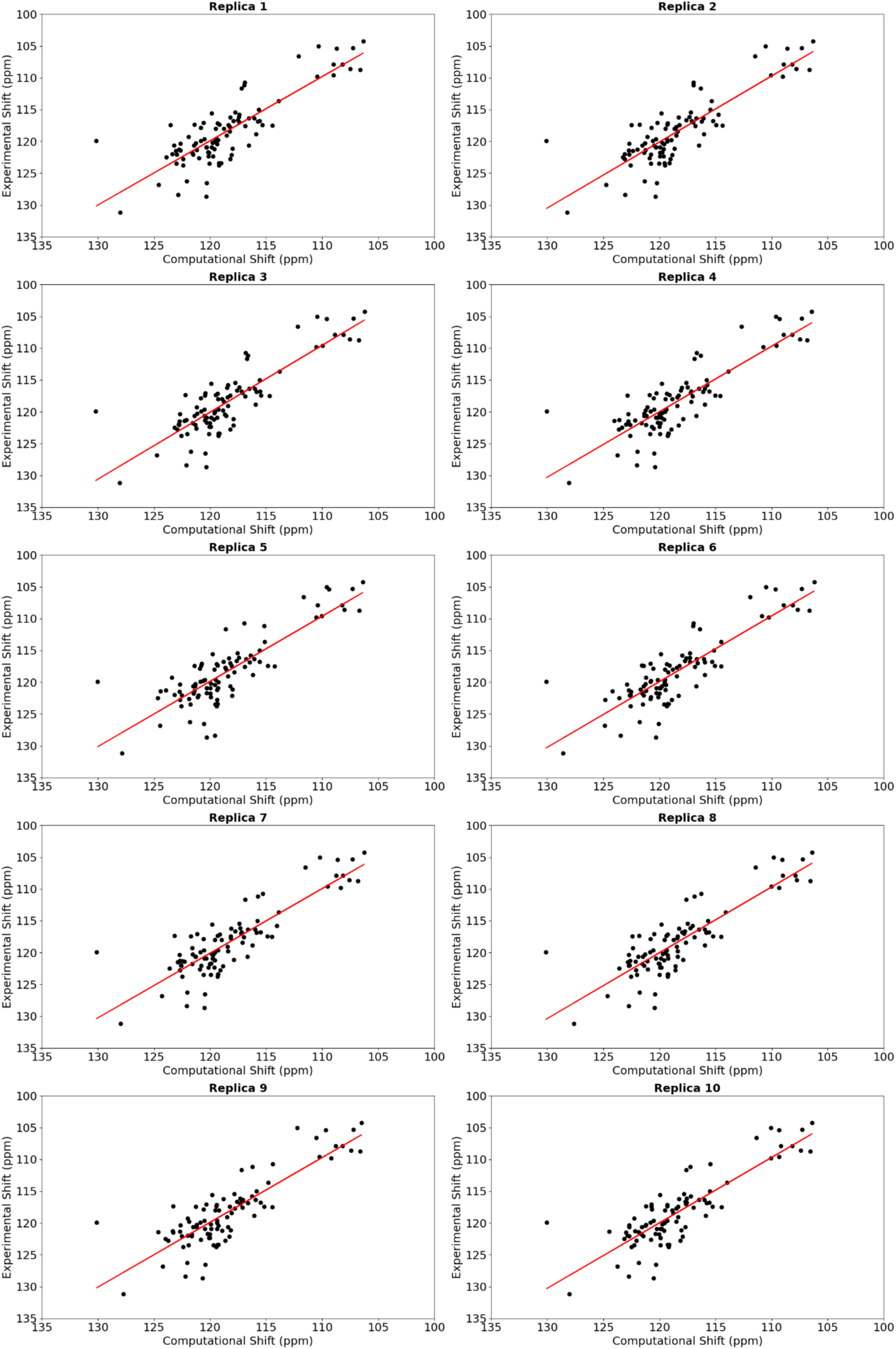
Chemical shift correlation for the N^H^ atom type between experimental and computational predictions for the neurotensin-1 receptor.

**Supplemental Figure 6.**
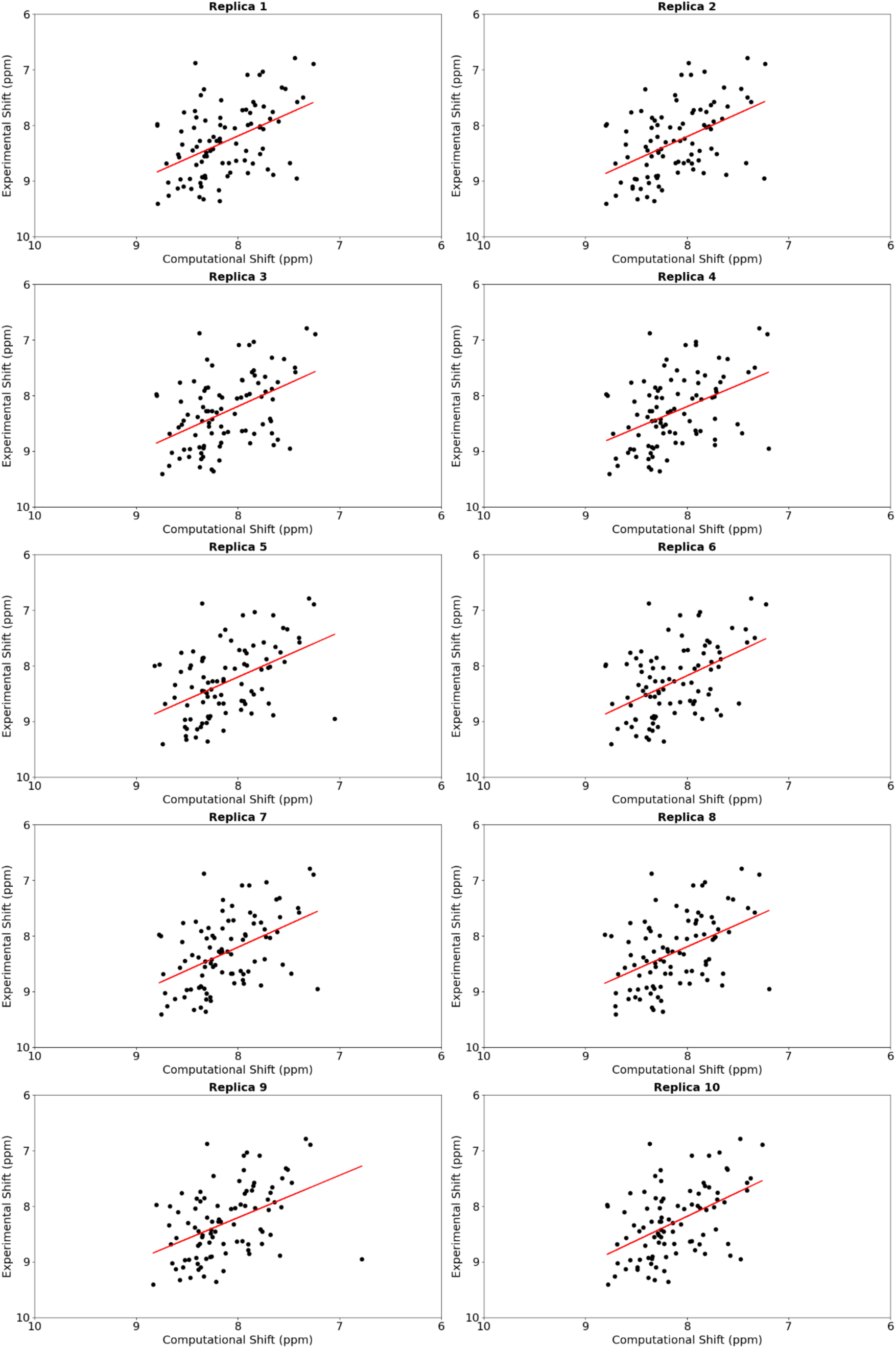
Chemical shift correlation for the H^N^ atom type between experimental and computational predictions for the neurotensin-1 receptor.

**Supplemental Figure 7.**
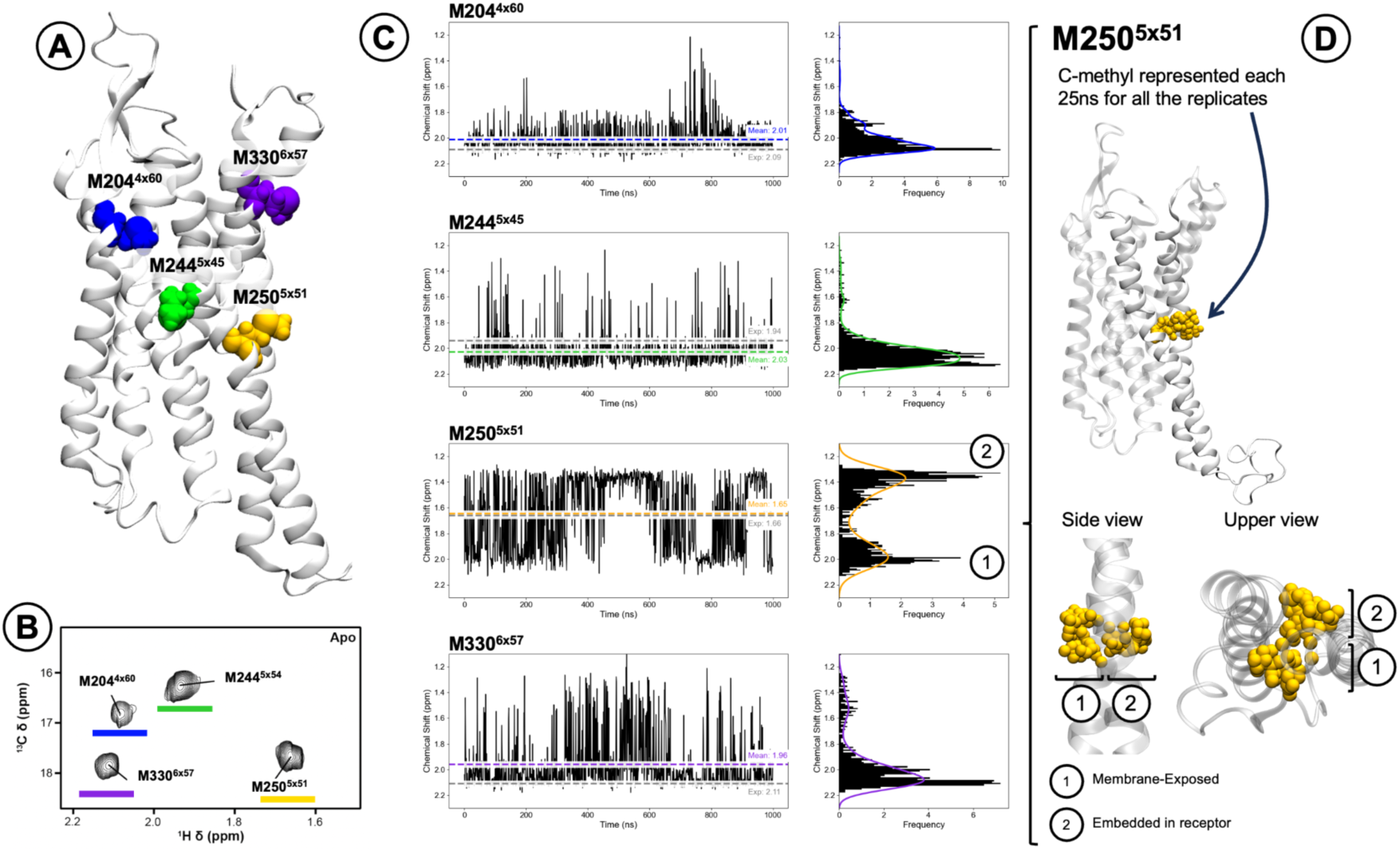
1H chemical shift profile of M204^4x60^, M244^5x45^, M250^5x51^ and M330^6x57^ in the neurotensin receptor 1. (**A**) Structural position of the selected methionines. (**B**) Experimental CS data for M250^5x51^ and M330^6x57^ extracted from the study of Bumbak *et al.* ^19^, **Figure 2**). (**C**) Predicted chemical shift, shown using the streaming tool. We highlighted the mean value from the predicted and experimental CS. An overlap between the predicted and the experimental chemical shift values can be seen for each of the methionines. (**D**) M250^5x51^ presents two different side chain conformations (yellow spheres) where the methyl group can be membrane-exposed or embedded in the receptor.

**Supplemental Figure 8.**
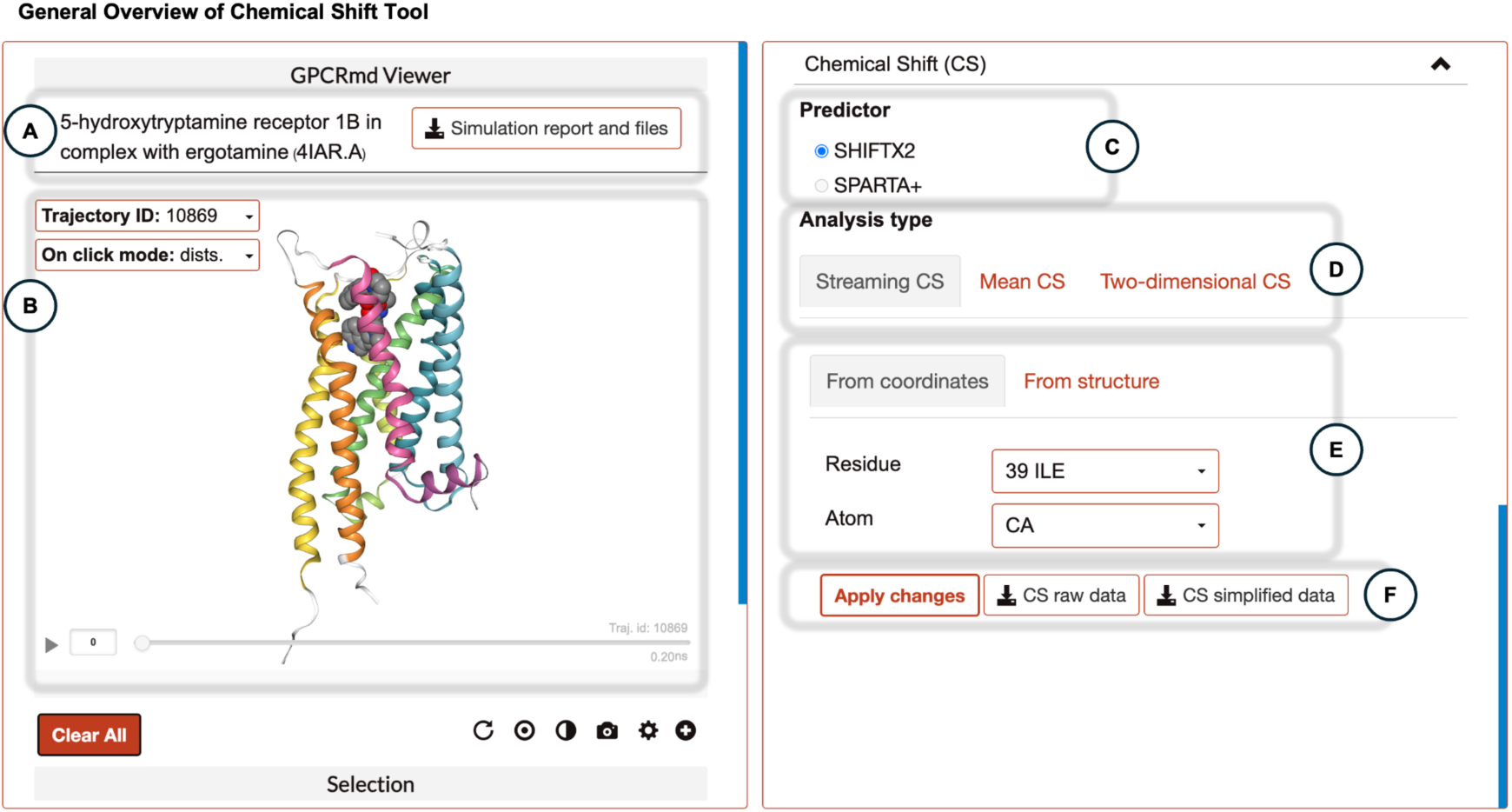
General overview of GPCRmd chemical shift tool. (**A**) Name of the receptor and PDB code. The simulation report and files button redirects the user to an in-depth page where all the necessary files to reproduce the MD and the trajectories can be downloaded. (**B**) GPCRmd viewer, where the user can visualize the MD simulation. (**C**) Allows to select which predictor to use when showing the results. (**D**) Allows to select a visualization mode to display the CS values. (**E**) Allows to select the atoms to display the chemical shifts. (**F**) Apply changes button will generate the plot with the specified visualization mode and atoms. The user can also download the csv files containing the chemical shifts for each atom on the system.

**Supplemental Figure 9.**
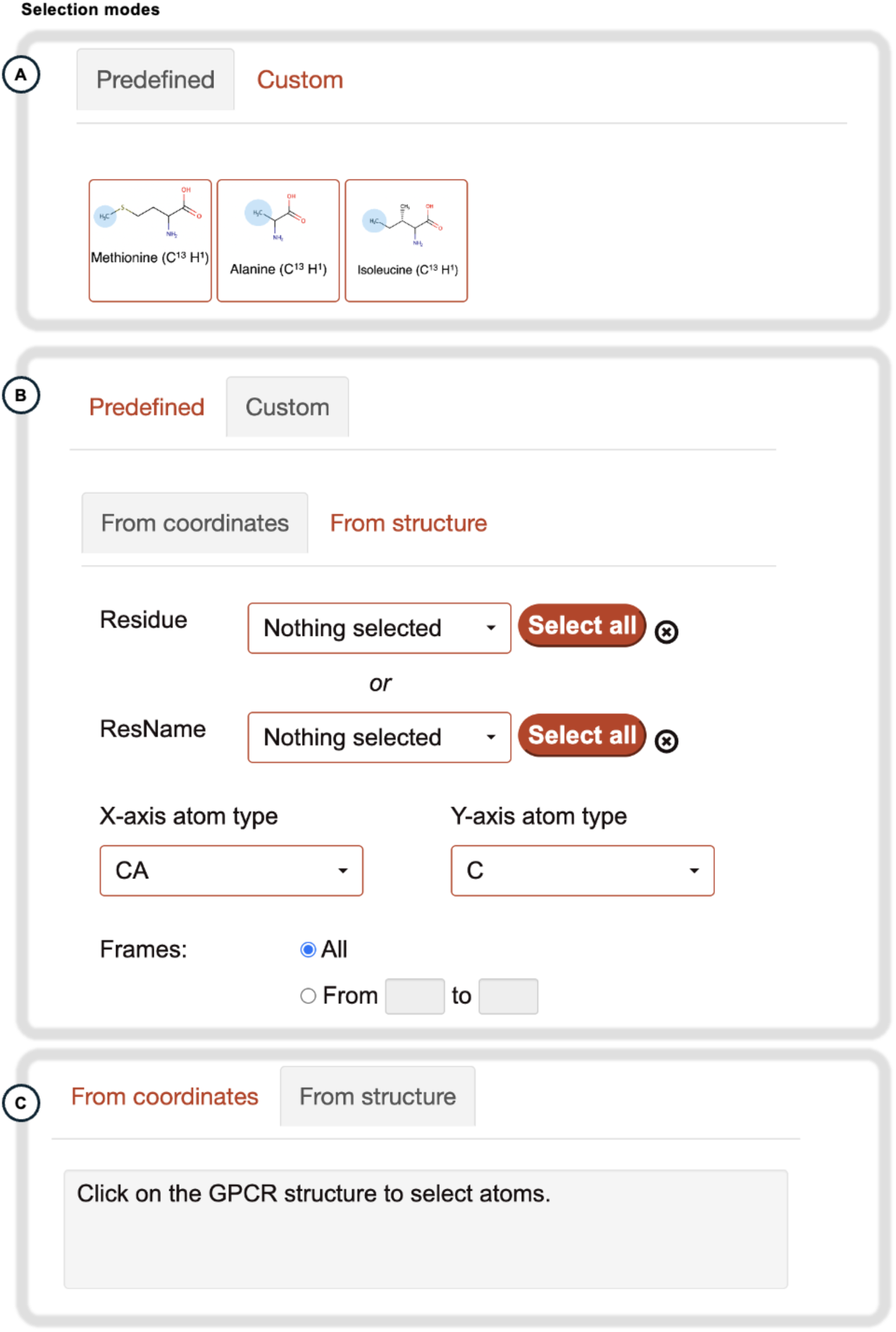
Selection modes in GPCRmd Chemical Shift tool. (**A**) There are 3 predefined groups of atoms based on the most common labeled atoms in NMR experiments (Methionines: Cε and Hε; Alanines: Cβ and Hβ; Isoleucines: Cδ and Hδ). This selection mode is not available in the streaming chemical shift mode because only 1 atom should be selected. (**B**) The user can choose between selecting residues by their IDs or names, as well as which atom types to include. This selection mode is specific for each visualization mode. (**C**) The user can also select atoms directly through the GPCRmd viewer

**Supplemental Figure 10.**
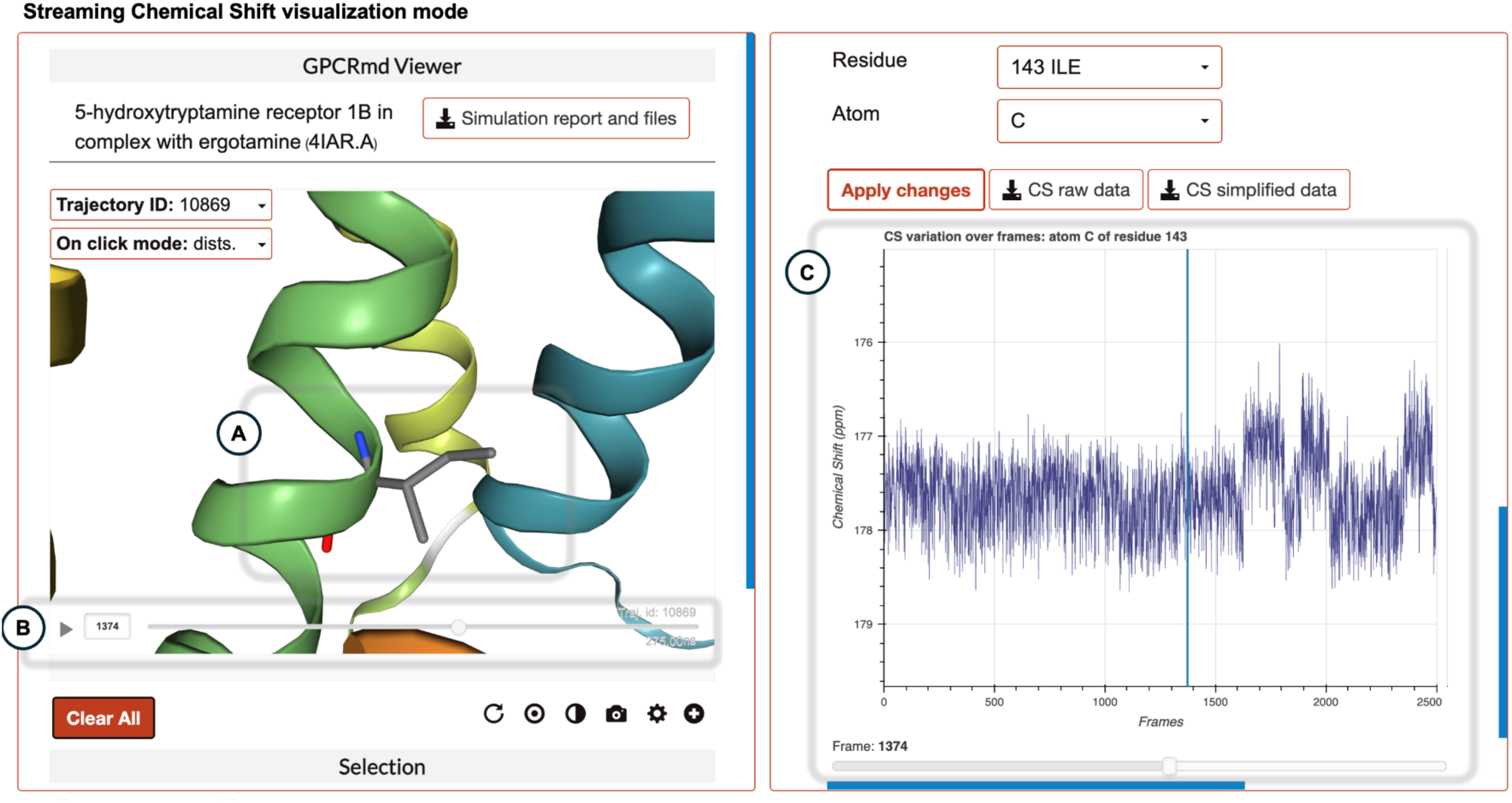
Streaming Chemical Shift visualization mode. In this mode the user can interactively see the evolution of the chemical shifts over the molecular simulation. (**A**) The selected atom is zoomed in as the CS values are displayed. (**B**) The user can play the molecular simulation using the GPCRmd viewer. (**C**) Evolution of the chemical shift for the selected residue. A blue line and a progress button keep track of the current frame displayed on the GPCRmd viewer. An accumulative histogram of the chemical shift values appears on the right part of the plot if the user hovers through it.

**Supplemental Figure 11.**
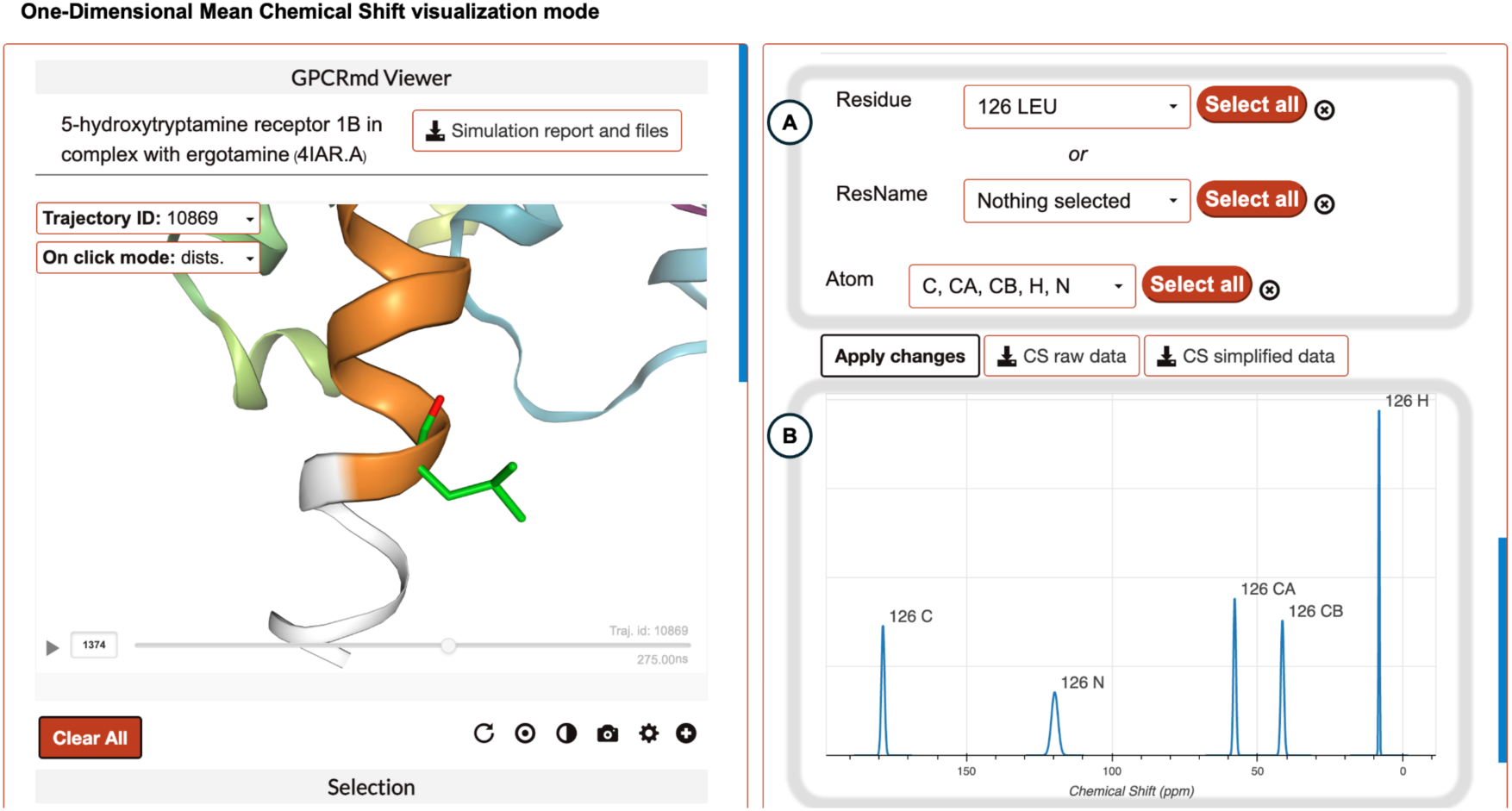
One-Dimensional Mean Chemical Shift visualization mode. This mode allows the user to quickly compare chemical shift values over multiple residues and atom types. (**A**) The selection mode allows the user to select multiple atom types and residues at the same time. (**B**) The mean signals for each selected atom are displayed. The width of the signal represents the associated error to the predicted value.

**Supplemental Figure 12.**
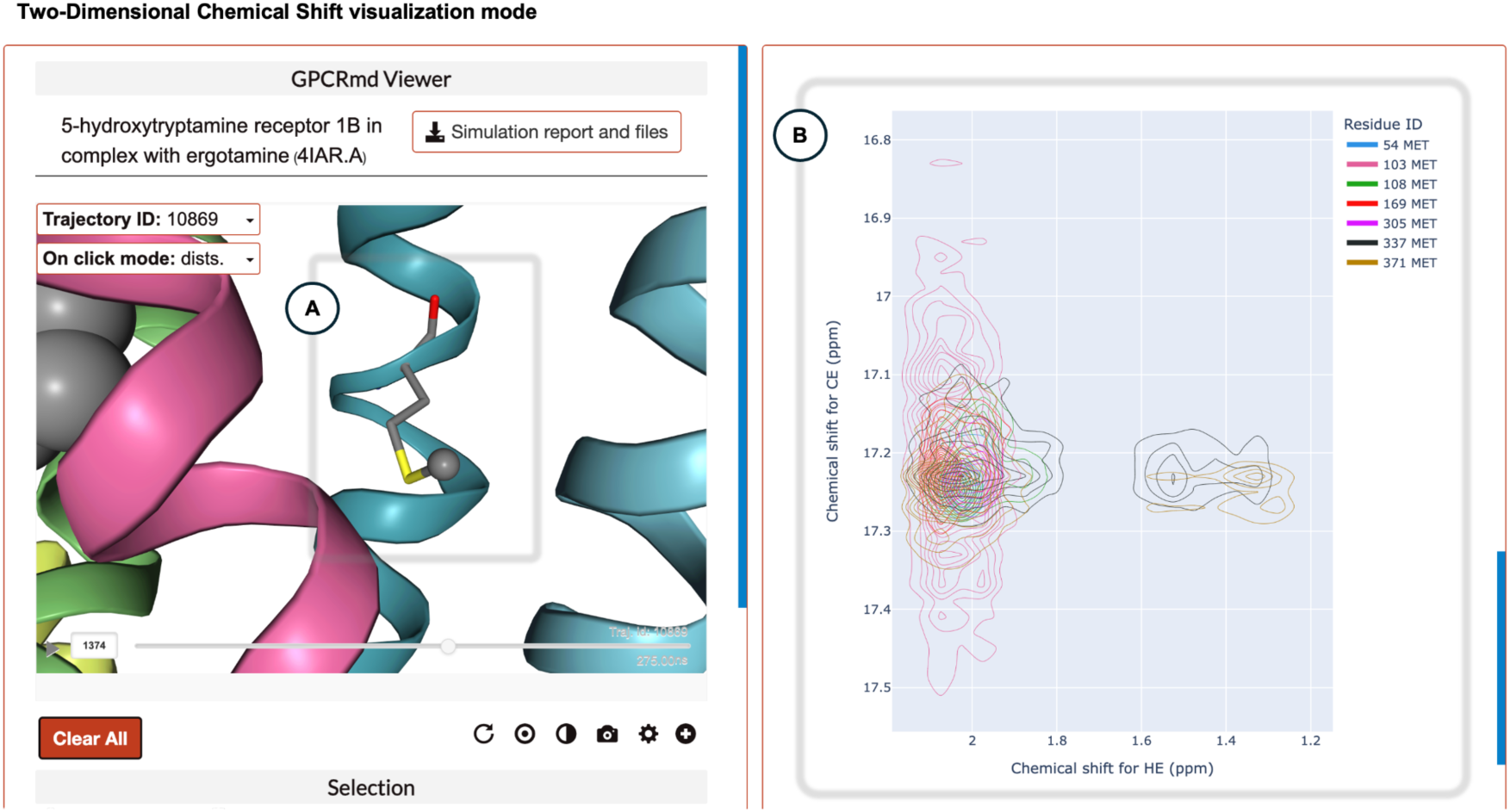
Two-Dimensional Chemical Shift visualization mode. This mode allows the user to compare different signals for 2 atoms over the same residues. It becomes especially useful when trying to detect which residues are the most different between the selected ones. (**A**) GPCRmd viewers zooms in on the selected residue in the right plot. (**B**) The user can interactively select the residues using the legend to get an isolated representation of their signal in the plot and it will be zoomed in the GPCRmd viewer.

## Bibliography

1. Lorente, J. S. et al. GPCR drug discovery: new agents, targets and indications. Nat. Rev. Drug Discov. (2025) doi:10.1038/s41573-025-01139-y.

2. García-Nafría, J. & Tate, C. G. Structure determination of GPCRs: cryo-EM compared with X-ray crystallography. Biochem. Soc. Trans. 49, 2345–2355 (2021).

3. Gusach, A., García-Nafría, J. & Tate, C. G. New insights into GPCR coupling and dimerisation from cryo-EM structures. Curr. Opin. Struct. Biol. 80, 102574 (2023).

4. Lopez-Balastegui, M. et al. Relevance of G protein-coupled receptor (GPCR) dynamics for receptor activation, signalling bias and allosteric modulation. Br. J. Pharmacol. (2024) doi:10.1111/bph.16495.

5. Schanda, P. & Haran, G. NMR and single-molecule FRET insights into fast protein motions and their relation to function. Annu. Rev. Biophys. 53, 247–273 (2024).

6. Hofmann, L. & Ruthstein, S. EPR spectroscopy provides new insights into complex biological reaction mechanisms. J. Phys. Chem. B 126, 7486–7494 (2022).

7. Kjær, V. M. S. et al. Ligand entry pathways control the chemical space recognized by GPR183. Chem. Sci. 14, 10671–10683 (2023).

8. Manglik, A. et al. Structural insights into the dynamic process of β2-adrenergic receptor signaling. Cell 161, 1101–1111 (2015).

9. Li, Y., Sun, J., Li, D. & Lin, J. The full activation mechanism of the adenosine A1 receptor revealed by GaMD and Su-GaMD simulations. Proc. Natl. Acad. Sci. U. S. A. 119, e2203702119 (2022).

10. Dwivedi-Agnihotri, H. et al. Distinct phosphorylation sites in a prototypical GPCR differently orchestrate β-arrestin interaction, trafficking, and signaling. Sci Adv 6, (2020).

11. Calebiro, D., Koszegi, Z., Lanoiselée, Y., Miljus, T. & O’Brien, S. G protein-coupled receptor-G protein interactions: a single-molecule perspective. Physiol. Rev. 101, 857–906 (2021).

12. Batebi, H. et al. Mechanistic insights into G-protein coupling with an agonist-bound G-protein-coupled receptor. Nat. Struct. Mol. Biol. 31, 1692–1701 (2024).

13. Kumar, G. A. et al. A molecular sensor for cholesterol in the human serotonin1A receptor. Sci Adv 7, (2021).

14. Thakur, N. et al. Anionic phospholipids control mechanisms of GPCR-G protein recognition. Nat. Commun. 14, 794 (2023).

15. Rodríguez-Espigares, I. et al. GPCRmd uncovers the dynamics of the 3D-GPCRome. Nat. Methods 17, 777–787 (2020).

16. Aranda-García, D. et al. Large scale investigation of GPCR molecular dynamics data uncovers allosteric sites and lateral gateways. Nat. Commun. 16, 2020 (2025).

17. Puthenveetil, R. & Vinogradova, O. Solution NMR: A powerful tool for structural and functional studies of membrane proteins in reconstituted environments. J. Biol. Chem. 294, 15914–15931 (2019).

18. Kimata, N., Reeves, P. J. & Smith, S. O. Uncovering the triggers for GPCR activation using solid-state NMR spectroscopy. J. Magn. Reson. 253, 111–118 (2015).

19. Mandala, V. S., Williams, J. K. & Hong, M. Structure and dynamics of membrane proteins from solid-state NMR. Annu. Rev. Biophys. 47, 201–222 (2018).

20. Jonas, E., Kuhn, S. & Schlörer, N. Prediction of chemical shift in NMR: A review. Magn. Reson. Chem. 60, 1021–1031 (2022).

21. Han, B., Liu, Y., Ginzinger, S. W. & Wishart, D. S. SHIFTX2: significantly improved protein chemical shift prediction. J. Biomol. NMR 50, 43–57 (2011).

22. Shen, Y. & Bax, A. SPARTA+: a modest improvement in empirical NMR chemical shift prediction by means of an artificial neural network. J. Biomol. NMR 48, 13–22 (2010).

23. Yin, Y.-L. et al. Molecular basis for kinin selectivity and activation of the human bradykinin receptors. Nat. Struct. Mol. Biol. 28, 755–761 (2021).

24. Joedicke, L. et al. The molecular basis of subtype selectivity of human kinin G-protein-coupled receptors. Nat. Chem. Biol. 14, 284–290 (2018).

25. Mohamadi, M., Goricanec, D., Wagner, G. & Hagn, F. NMR sample optimization and backbone assignment of a stabilized neurotensin receptor. J. Struct. Biol. 215, 107970 (2023).

26. Dixon, A. D. et al. Effect of ligands and transducers on the neurotensin receptor 1 conformational ensemble. J. Am. Chem. Soc. 144, 10241–10250 (2022).

27. Bender, A. M., Parr, L. C., Livingston, W. B., Lindsley, C. W. & Merryman, W. D. 2B determined: The future of the serotonin receptor 2B in drug discovery. J. Med. Chem. 66, 11027–11039 (2023).

28. Wang, C. et al. Structural basis for molecular recognition at serotonin receptors. Science 340, 610–614 (2013).

29. Bumbak, F. et al. Ligands selectively tune the local and global motions of neurotensin receptor 1 (NTS1). Cell Rep. 42, 112015 (2023).

30. Yang, A., Yu, G., Wu, Y. & Wang, H. Role of β2-adrenergic receptors in chronic obstructive pulmonary disease. Life Sci. 265, 118864 (2021).

31. Scharf, M. M. et al. A focus on unusual ECL2 interactions yields β2 -adrenergic receptor antagonists with unprecedented scaffolds. ChemMedChem 15, 882–890 (2020).

32. Koehl, A. et al. Structure of the µ-opioid receptor-Gi protein complex. Nature 558, 547– 552 (2018).

33. Kapoor, A., Martinez-Rosell, G., Provasi, D., de Fabritiis, G. & Filizola, M. Dynamic and kinetic elements of µ-opioid receptor functional selectivity. Sci. Rep. 7, 11255 (2017).

34. Hofmann, K. P. & Lamb, T. D. Rhodopsin, light-sensor of vision. Prog. Retin. Eye Res. 93, 101116 (2023).

35. Rose, A. S. & Hildebrand, P. W. NGL Viewer: a web application for molecular visualization. Nucleic Acids Res. 43, W576–9 (2015).

36. Rose, A. S. et al. NGL viewer: web-based molecular graphics for large complexes. Bioinformatics 34, 3755–3758 (2018).

37. Humphrey, W., Dalke, A. & Schulten, K. VMD: visual molecular dynamics. J. Mol. Graph. 14, 33–8, 27–8 (1996).

