## Supplementary Material for "Integrative Biophysical Illumination of the 3D GPCRome Dynamics"

**Supplementary Note 1:**
The first mode, termed "One-Dimensional Mean Chemical Shift", presents a mean representation of the CS derived from the entire MD. The distribution's center corresponds to the mean of the CS, while its width represents an error estimation for the CS prediction (refer to methods for more details). This error estimation guides the user to distinguish whether two measures are significantly different or not. In **Figure 4B**, a visual representation illustrates how the two distributions for the Cα and Nitrogen atoms between the aspartates differ in CS. For instance, the Nitrogen atom and the Cα of $D79^{2.50}$ and $D192^{45x51}$ are well separated in the $\beta_2$ adrenergic receptor. The second mode, termed "Two-Dimensional Chemical Shift", allows users to correlate two atom types by mapping their distribution on a plane, resembling a real 2D-NMR plot. This mode helps to quickly understand the ranges of CS values explored throughout the simulation. **Figure 4C** illustrates the "Two-Dimensional Chemical Shift" mode, where $D79^{2.50}$ has slightly greater fluctuation in CS compared to $D192^{45x51}$ for the Cα and Nitrogen atoms. Importantly, both visualization modes are interactive to enhance user comprehension. For instance, selecting a specific NMR signal allows users to visualize the corresponding atom within the 3D receptor structure using the GPCRmd viewer. A snapshot from GPCRmd highlighting the main features of both modes is presented in **Figures S8-S12**.

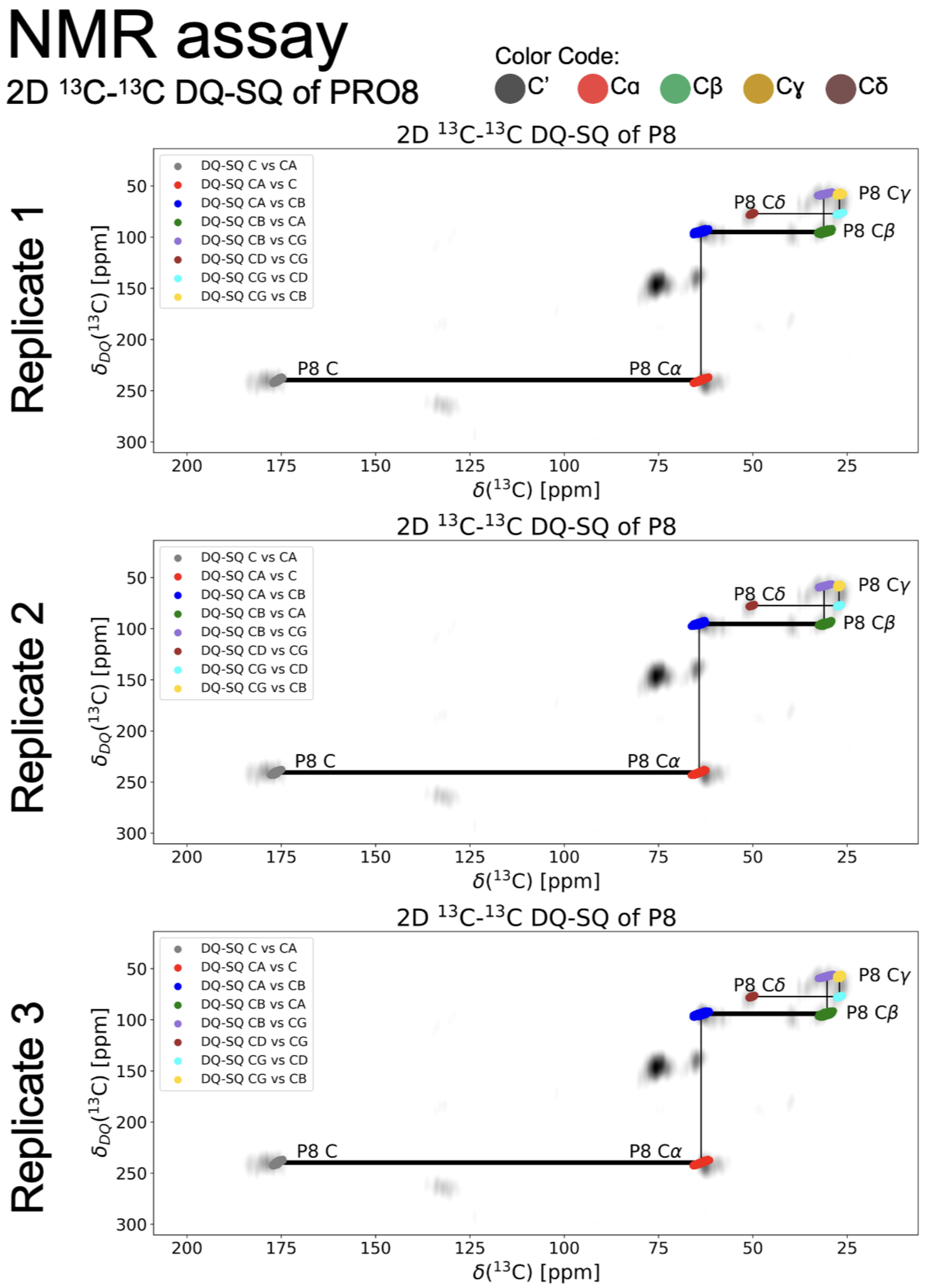


**Supplemental Figure 1. $^{13}C$-$^{13}C$ DQ-SQ overlap between predicted (colored) and experimental (grey) chemical shift values for P8 in the Bradykinin 1 Receptor.** The lines between atom types represent the carbon-carbon correlations, indicating spatial proximity between them. Each replicate consists in a independent MD simulation of 500ns (GPCRmd ID: 2095, PDB: 7EIB).

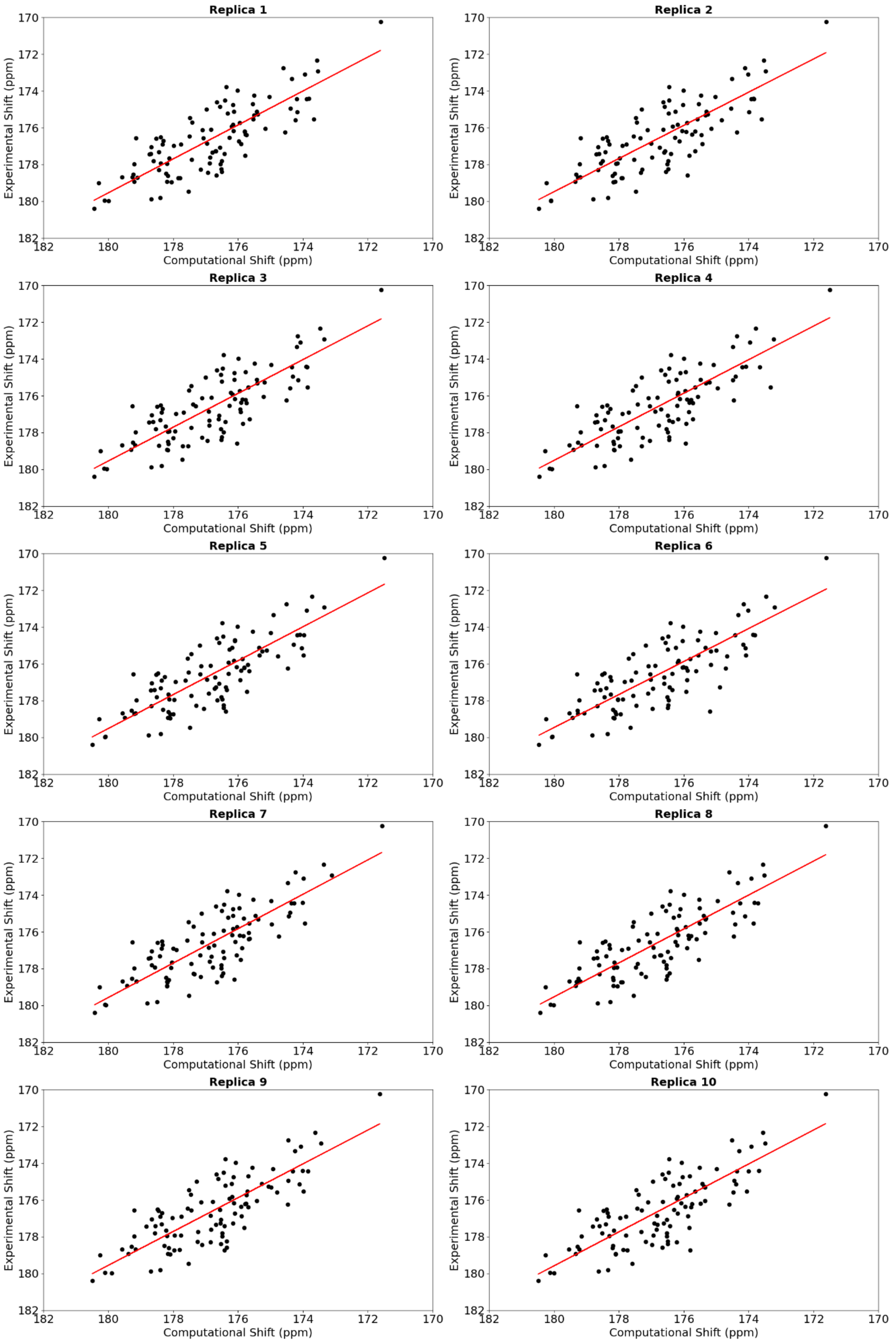


**Supplemental Figure 2. Chemical shift correlation for the C atom type between experimental and computational predictions for the neurotensin-1 receptor**

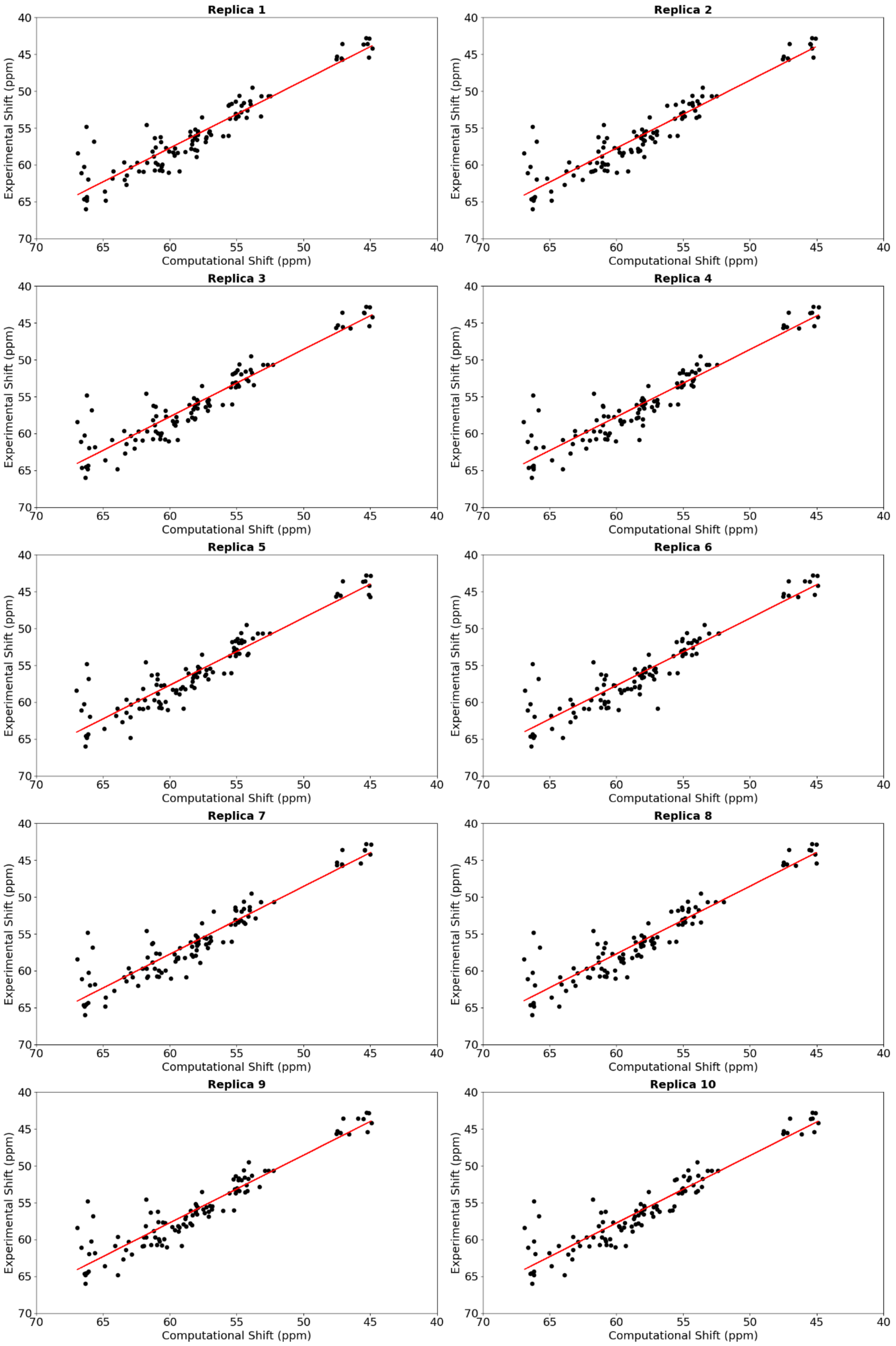


**Supplemental Figure 3. Chemical shift correlation for the Cα atom type between experimental and computational predictions for the neurotensin-1 receptor**

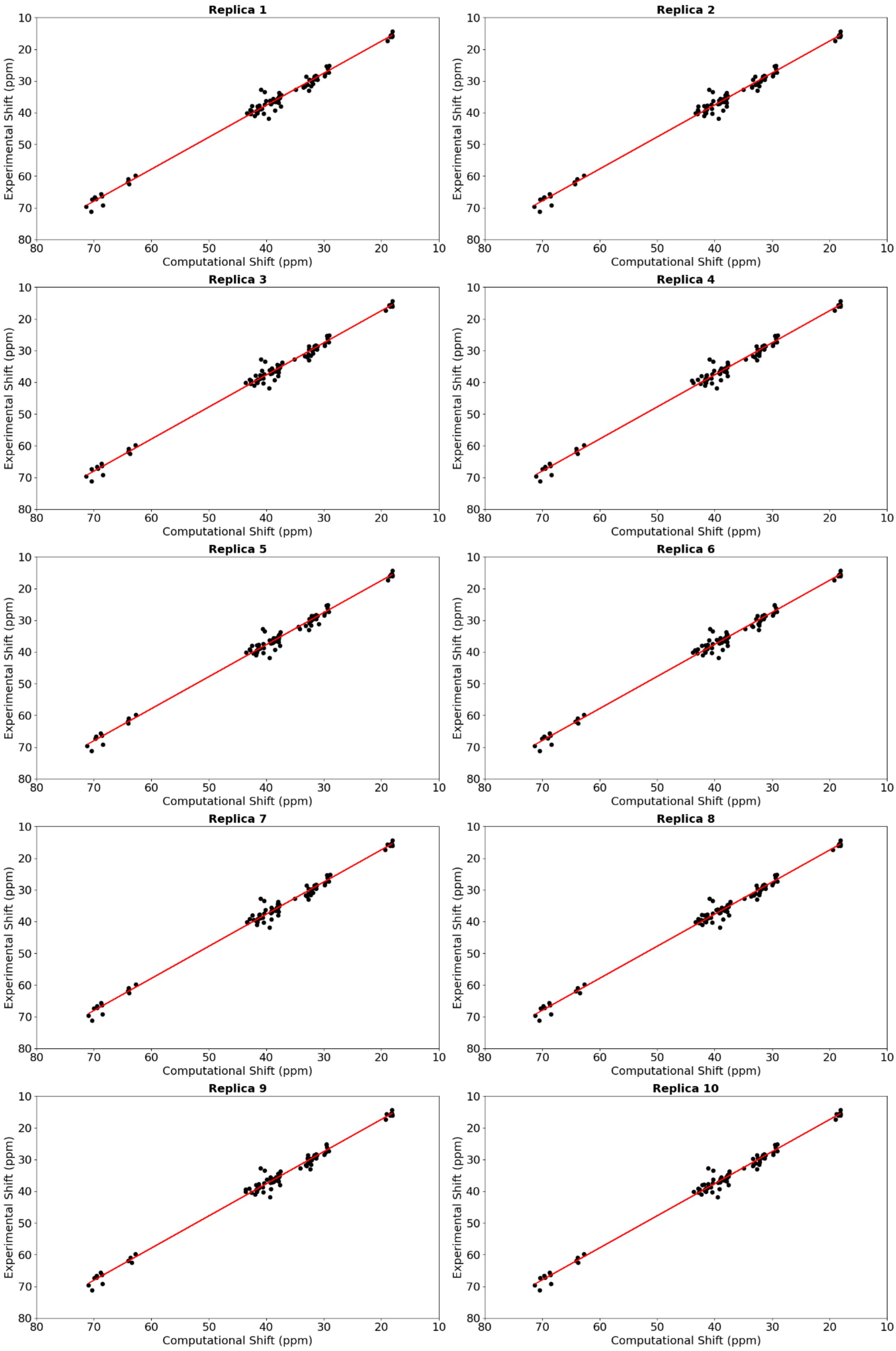


**Supplemental Figure 4. Chemical shift correlation for the Cβ atom type between experimental and computational predictions for the neurotensin-1 receptor**

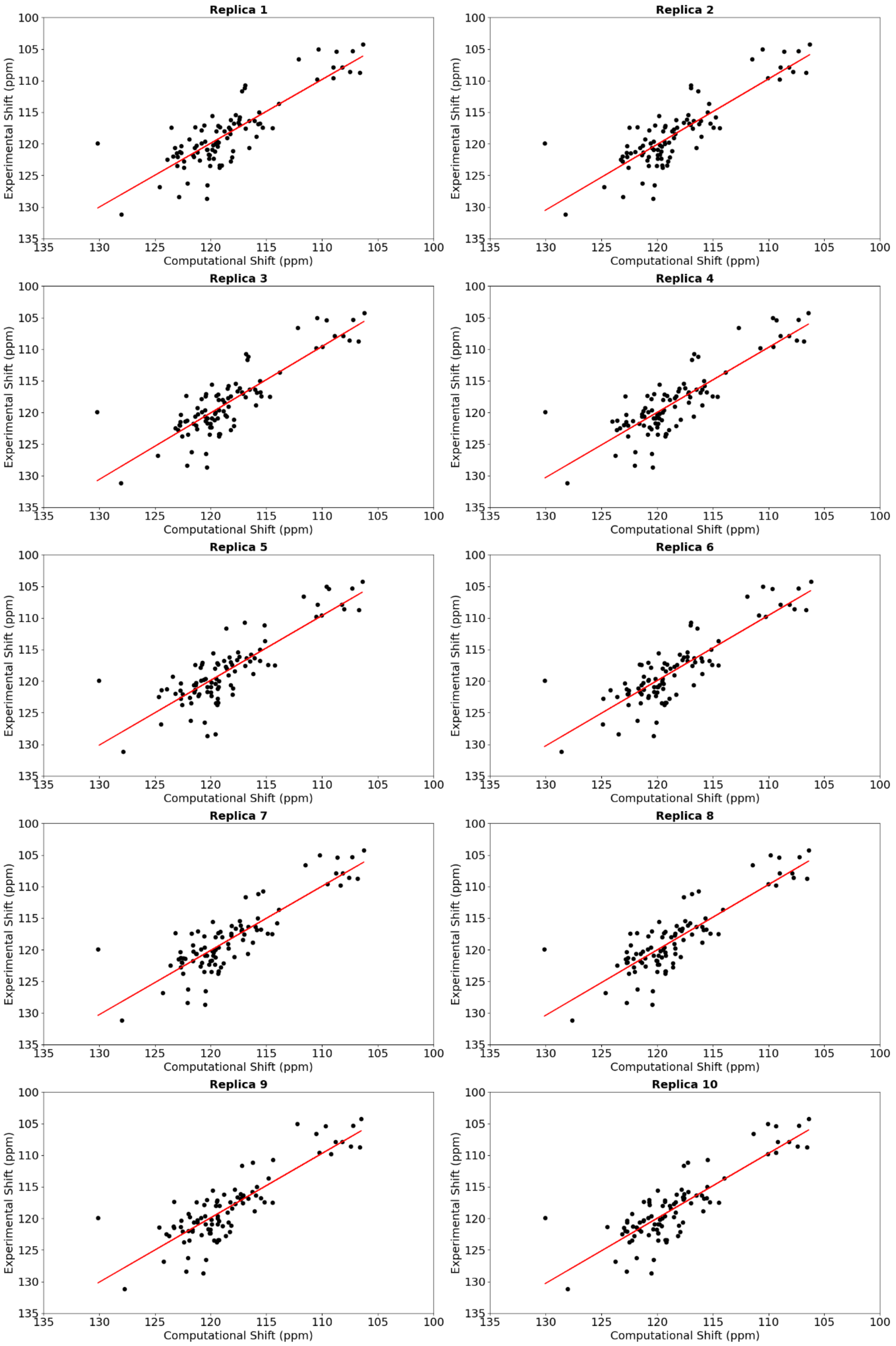


**Supplemental Figure 5. Chemical shift correlation for the $N^H$ atom type between experimental and computational predictions for the neurotensin-1 receptor**

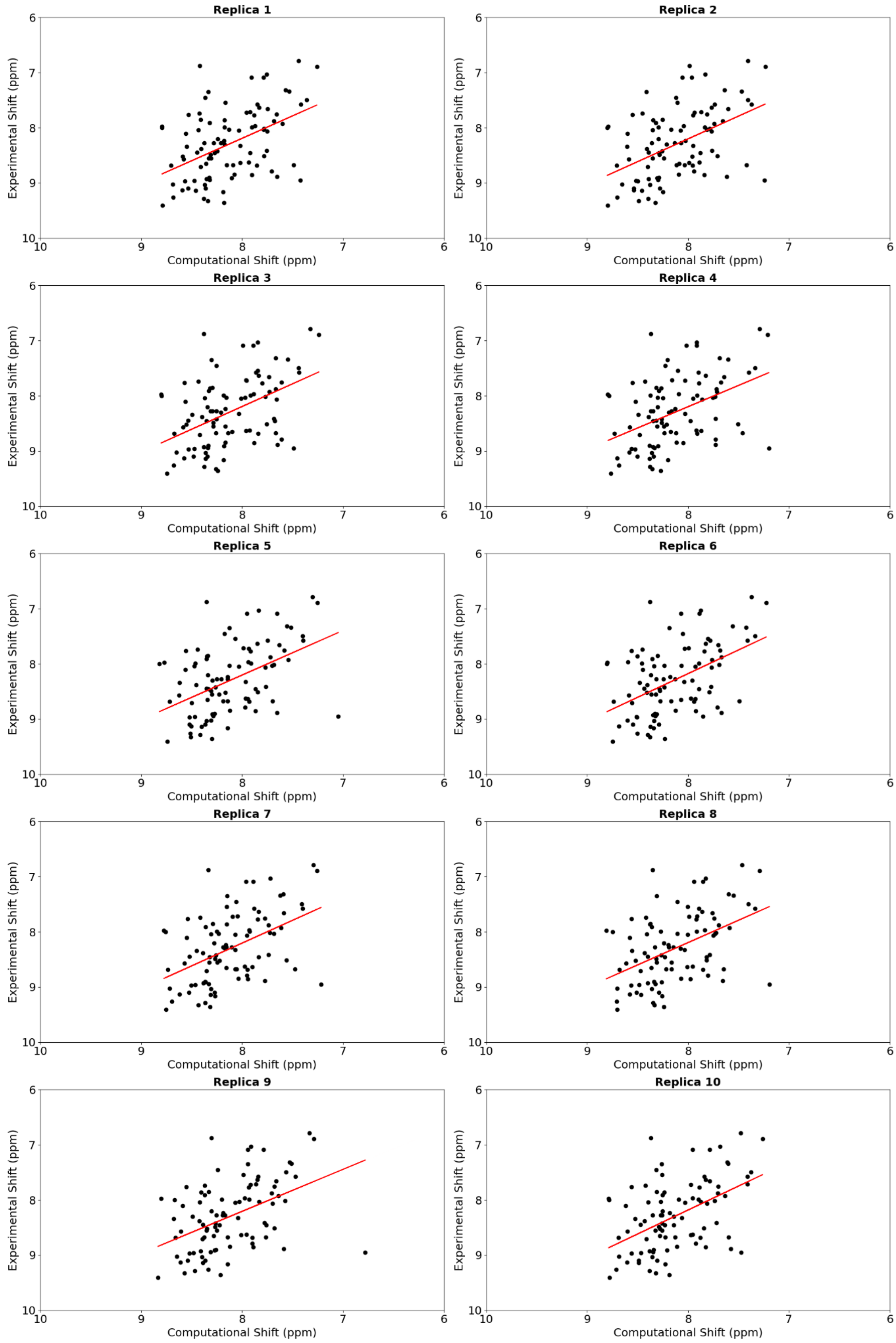


**Supplemental Figure 6. Chemical shift correlation for the $H^N$ atom type between experimental and computational predictions for the neurotensin-1 receptor**

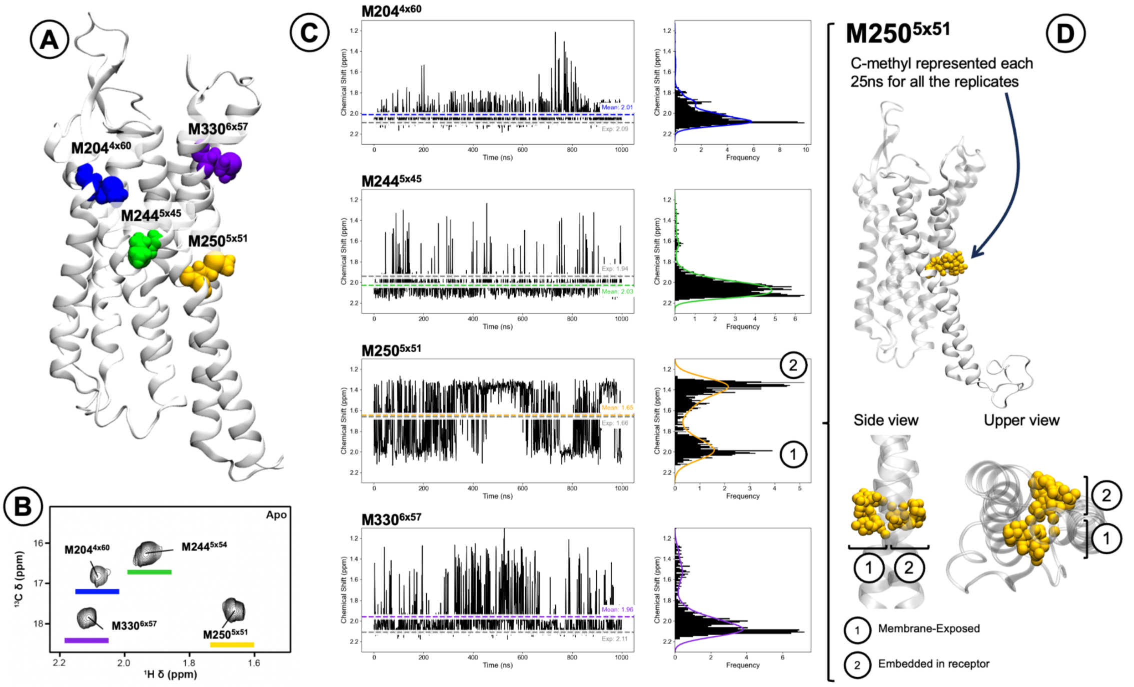


**Supplemental Figure 7. $^1$H chemical shift profile of M204$^{4x60}$, M244$^{5x45}$, M250$^{5x51}$ and M330$^{6x57}$ in the neurotensin receptor 1.** (**A**) Structural position of the selected methionines. (**B**) Experimental CS data for M250$^{5x51}$ and M330$^{6x57}$ extracted from the study of Bumbak *et al.* [19], **Figure 2**). (**C**) Predicted chemical shift, shown using the streaming tool. We highlighted the mean value from the predicted and experimental CS. An overlap between the predicted and the experimental chemical shift values can be seen for each of the methionines. (**D**) M250$^{5x51}$ presents two different side chain conformations (yellow spheres) where the methyl group can be membrane-exposed or embedded in the receptor.

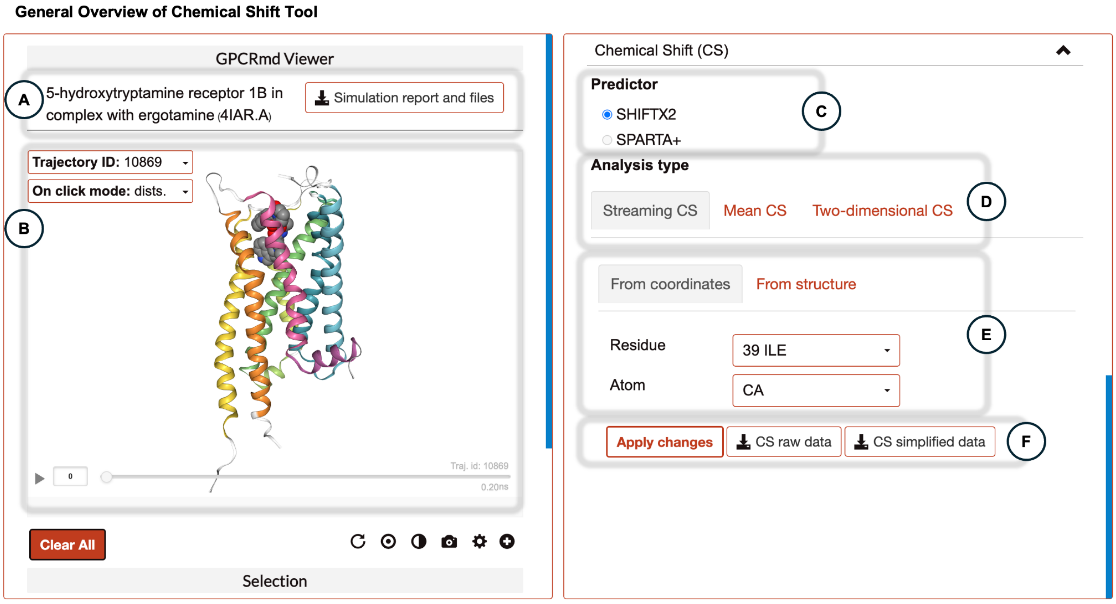


**Supplemental Figure 8. General overview of GPCRmd chemical shift tool.** (**A**) Name of the receptor and PDB code. The simulation report and files button redirects the user to an in-depth page where all the necessary files to reproduce the MD and the trajectories can be downloaded. (**B**) GPCRmd viewer, where the user can visualize the MD simulation. (**C**) Allows to select which predictor to use when showing the results. (**D**) Allows to select a visualization mode to display the CS values. (**E**) Allows to select the atoms to display the chemical shifts. (**F**) Apply changes button will generate the plot with the specified visualization mode and atoms. The user can also download the csv files containing the chemical shifts for each atom on the system.

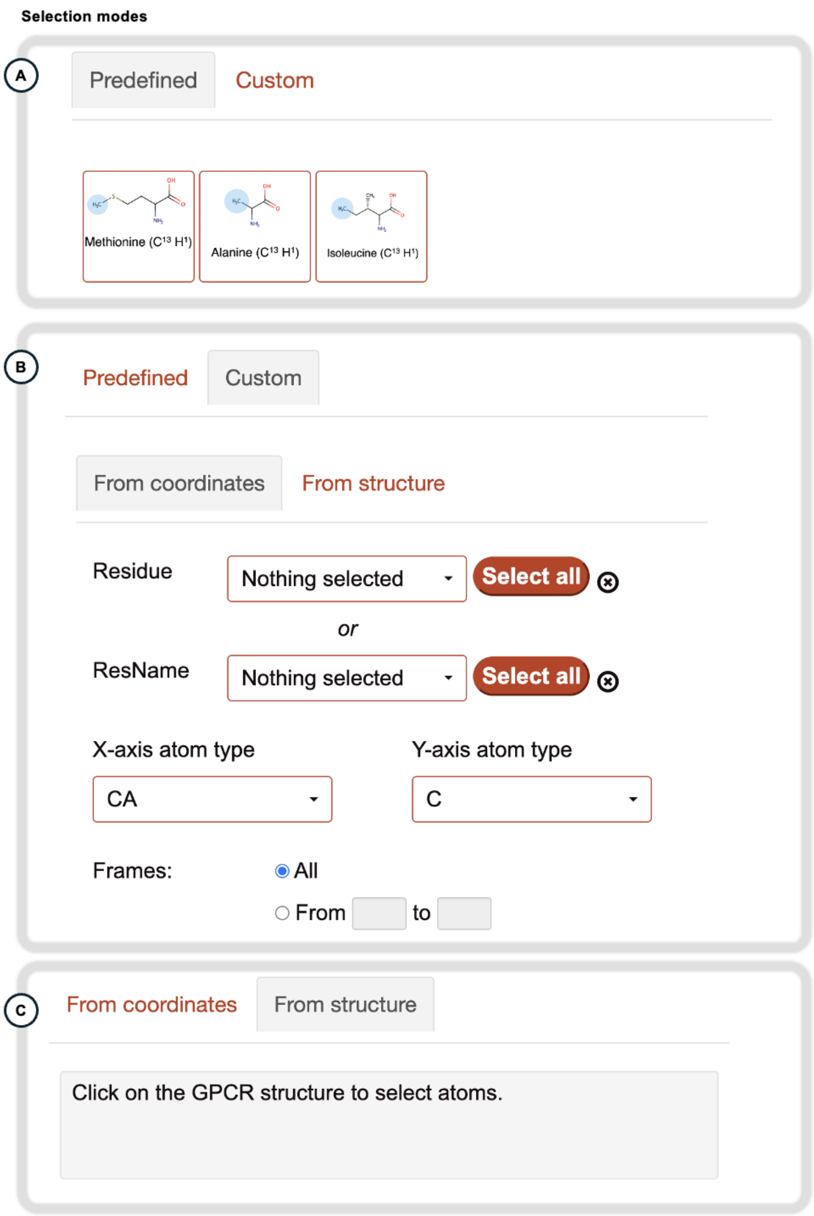


**Supplemental Figure 9. Selection modes in GPCRmd Chemical Shift tool.** (**A**) There are 3 predefined groups of atoms based on the most common labeled atoms in NMR experiments (Methionines: Cε and Hε; Alanines: Cβ and Hβ; Isoleucines: Cδ and Hδ). This selection mode is not available in the streaming chemical shift mode because only 1 atom should be selected. (**B**) The user can choose between selecting residues by their IDs or names, as well as which atom types to include. This selection mode is specific for each visualization mode. (**C**) The user can also select atoms directly through the GPCRmd viewer

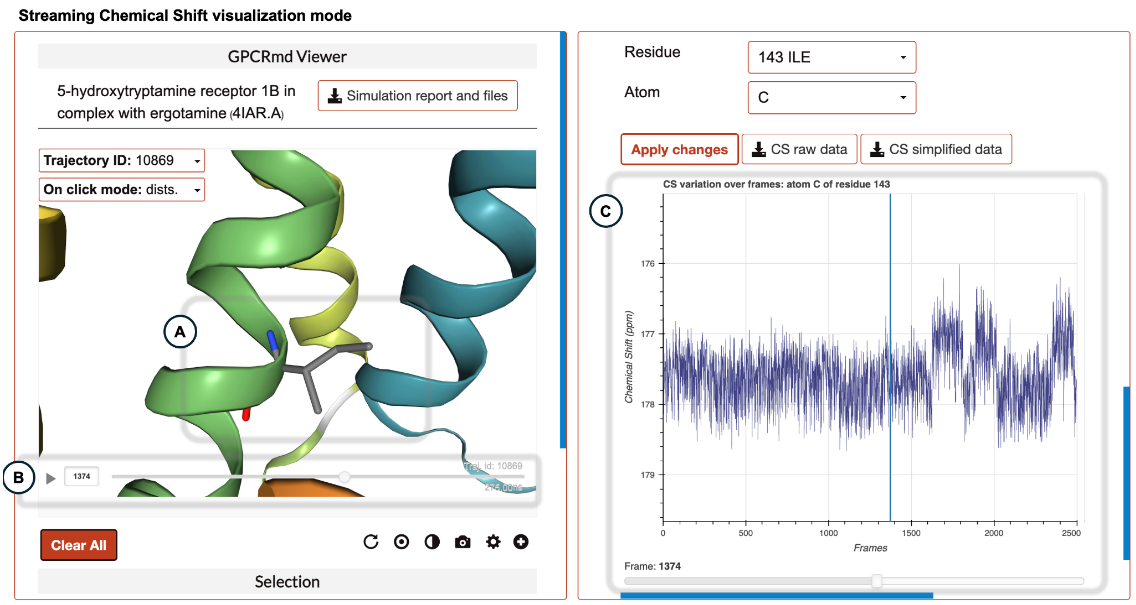


**Supplemental Figure 10. Streaming Chemical Shift visualization mode.** In this mode the user can interactively see the evolution of the chemical shifts over the molecular simulation. (**A**) The selected atom is zoomed in as the CS values are displayed. (**B**) The user can play the molecular simulation using the GPCRmd viewer. (**C**) Evolution of the chemical shift for the selected residue. A blue line and a progress button keep track of the current frame displayed on the GPCRmd viewer. An accumulative histogram of the chemical shift values appears on the right part of the plot if the user hovers through it.

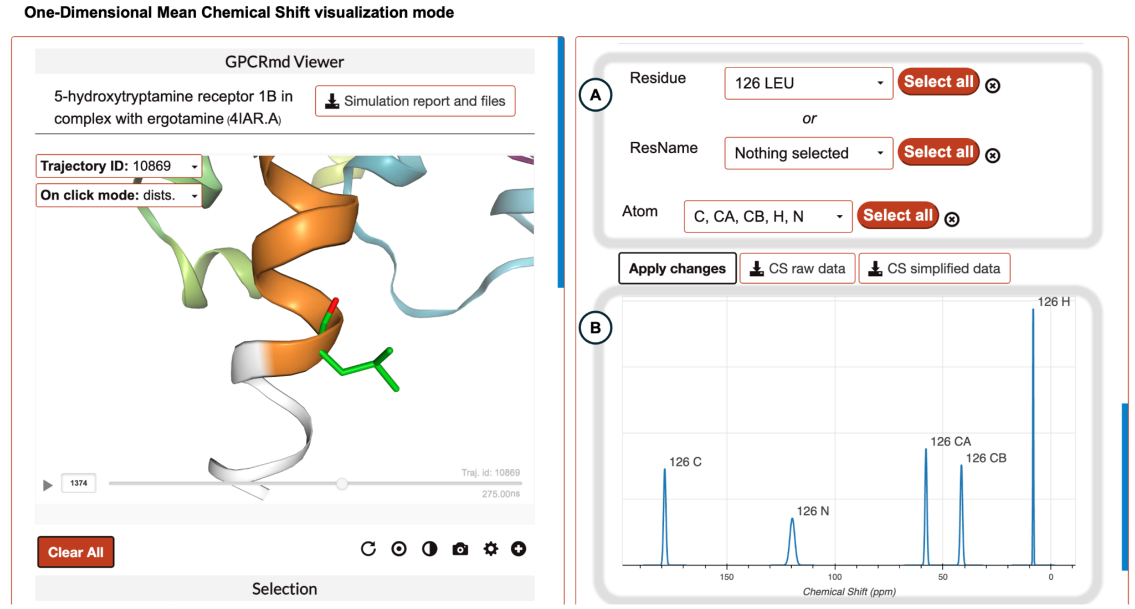


**Supplemental Figure 11. One-Dimensional Mean Chemical Shift visualization mode.** This mode allows the user to quickly compare chemical shift values over multiple residues and atom types. (**A**) The selection mode allows the user to select multiple atom types and residues at the same time. (**B**) The mean signals for each selected atom are displayed. The width of the signal represents the associated error to the predicted value.

**Two-Dimensional Chemical Shift visualization mode**

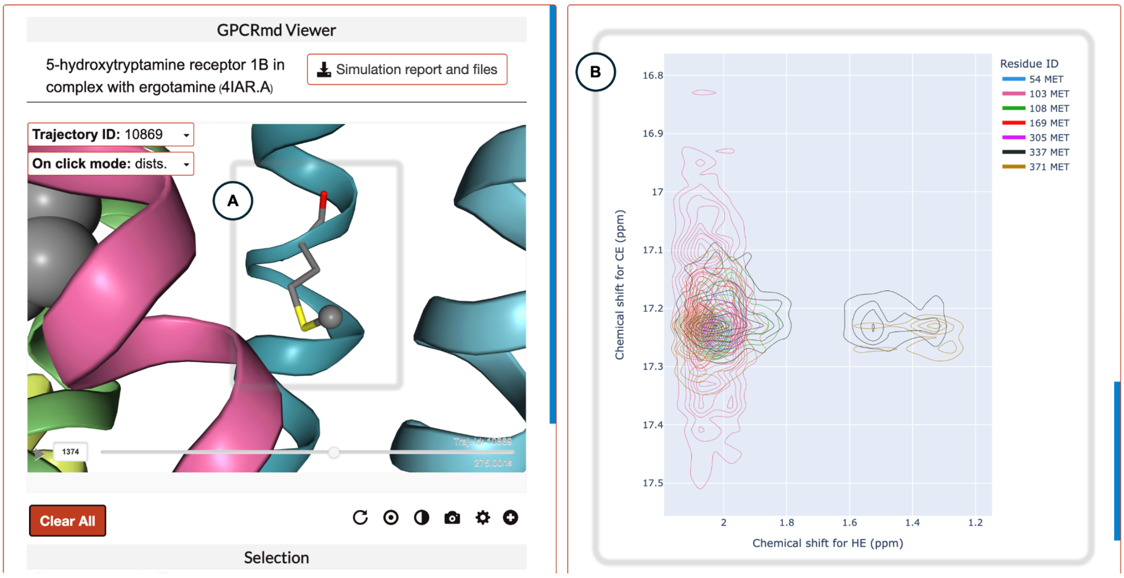


**Supplemental Figure 12. Two-Dimensional Chemical Shift visualization mode.** This mode allows the user to compare different signals for 2 atoms over the same residues. It becomes especially useful when trying to detect which residues are the most different between the selected ones. (**A**) GPCRmd viewers zooms in on the selected residue in the right plot. (**B**) The user can interactively select the residues using the legend to get an isolated representation of their signal in the plot and it will be zoomed in the GPCRmd viewer.
